# Comprehensive analysis of myeloid reporter mice

**DOI:** 10.1101/2025.02.24.639159

**Authors:** Yidan Wang, Samuel D. Dowling, Vanessa Rodriguez, Jessica Maciuch, Meghan Mayer, Tyler Therron, Tovah N. Shaw, Miranda G. Gurra, Caroline L. Shah, Hadijat-Kubura M. Makinde, Florent Ginhoux, David Voehringer, Cole A. Harrington, Toby Lawrence, John R. Grainger, Carla M. Cuda, Deborah R. Winter, Harris R. Perlman

## Abstract

Macrophages are a pivotal cell type within the synovial lining and sub-lining of the joint, playing a crucial role in maintaining homeostasis of synovium. Although fate-mapping techniques have been employed to differentiate synovial macrophages from other synovial myeloid cells, no comprehensive study has yet been conducted within the mouse synovial macrophage compartment. In this study, we present, for the first time, lineage tracing results from 18 myeloid-specific fate-mapping models in mouse peripheral blood (PB) and synovial tissue. The identification of synovial macrophages and monocyte-lineage cells through flow cytometry was further validated using cellular indexing of transcriptomes and epitopes by sequencing (CITE-seq) datasets. These findings provide a valuable methodological tool for researchers to select appropriate models for studying the function of synovial myeloid cells and serve as a reference for investigations in other tissue types.

## INTRODUCTION

Peripheral blood (PB) monocytes exist as at least 3 populations, classical monocytes (CM), intermediate (IM) and non-classical monocytes (NCM) ^1^. In many tissues, monocytes do not contribute to the macrophage population during steady-state conditions, although they do replenish macrophages in organs such as skin, colon, or lung ^2^. However, monocytes play a critical role during the response to insults by extravasating into tissue and differentiating into monocyte-derived macrophages ^1^. The synovial lining is crucial to maintain homeostasis and ensure the integrity of cartilage and bone. Macrophages and fibroblasts are the two cell types that comprise the synovial lining. Synovial tissue macrophages are embryonically derived and exist as a heterogenous population that is dependent on their local environment ^3–5^. We are the first to show macrophage heterogeneity in the synovium of mice and in patients with rheumatoid arthritis as well as in healthy controls ^6,7^. More recently we discovered a new population of tissue-resident monocyte-lineage cells, (TRMC) which are transcriptionally distinct from PB monocytes and tissue macrophages, derived from an embryonic origin and capable of self-repopulating in tissue^8^. Studies by Culemann *et al*., and Montgomery *et al*., utilized Cx3cr1^CreERT2^ mice to further distinguish macrophage and TRMC populations, respectively ^5,8^. Additionally, numerous investigators employed known macrophage-specific Cre mice to delete genes of interest such as Fas, Caspase 8, Flip, and Irf5 in the synovium ^9–23^. Yet, none of these studies determined the expression of macrophage specific Cre in synovial subpopulations. Further, while multiparameter and single-cell RNA sequencing (scRNA-seq) contribute to identifying individual populations, a current unmet need is the generation of lineage tracing models to accurately target a subpopulation. Fate-mapping is a method for tracing the origin of cell populations by labelling their progenitor cells. Monocyte-lineage and macrophage fate-mapping studies use fluorescent proteins down-stream of a cell type-specific promoter or express Cre recombinase which is then crossed with a loxP-stop-loxP reporter mouse. Over the past years several fate mapping and reporter mice such as LysM^Cre^, Csf1r^Cre^, Runx^Cre^, Tnfrsf11^Cre^, Cx3cr1^Cre^, Cx3cr1^CreERT2^, and more recently Ms4a3^Cre^ as well as Cx3cr1^GFP^, Ccr2^GFP^, and LysM^GFP^ knock-in mice have been extensively studied across mouse tissues to assess their efficacy in identifying the embryonic origin of monocytes ^2^. In the last few years, Tmem119^GFP^ ^24^ knock-in mice and P2ry12^CreERT2^ ^25^ mice have demonstrated efficient labeling capabilities for specific tissue macrophage populations, such as microglia, while Lyve1^Cre^ mice identifies perivascular macrophages ^26–29^. While these studies have been crucial for identifying whole populations of macrophages, none of these studies centered on designing fate mapping models to decipher macrophage heterogeneity in the synovial tissue.

In these studies, we accumulated 18 conditional or constitutive fate-mapping mice to provide a comprehensive map for myeloid lineage tracing in murine synovium. We validated these studies using cellular indexing of transcriptomes and epitopes by sequencing (CITE-seq) of the murine synovium. Together, our results will serve as a reference for future researchers to design models for specifically targeting synovial myeloid populations to investigate their ontogeny and functions.

## METHODS

### Generation of reporter mice

Breeder pairs of mice including Cx3cr1^GFP/GFP^ (005582), Cx3cr1^Cre/Cre^ (025524), Cx3cr1^CreERT2/CreERT2^ (020940), Ccr2^GFP/GFP^ (027619), Nur77 (Nr4a1)^RFP^ (034918), Csf1r^Cre^ (029206), hCD68^GFP^ (026827), LysM^Cre/+^ (004781), Lyve-1^GFP-Cre/+^ (012601), Pf4^iCre^ (008535), Ms4a3^Cre/+^ (036382 used for data generation; generously gifted from Florent Ginhoux for pilot studies ^30^), P2ry12^CreERT2/CreERT2^ (034727), Tmem119^GFP/GFP^ (031823), Mrp8^Cre-ires-GFP^ (021614), and Ai3^YFP/YFP^ (007903) were purchased from Jackson Laboratory and bred in specific pathogen-free animal facility at the Center for Comparative Medicine, Northwestern University. Cd74^tdTomato^ mice were a generous gift from Dr. Cole Harrington ^31^, Timd4^Cre/+^Rosa^YFP^ mice were a generous gift from Dr. John Grainger ^32^, Cd163^iCre^ mice were a generous gift from Dr. Toby Lawrence ^33^ and Retnla^Cre^ mice were a generous gift from David Voehringer ^34^. All mice used in this study were heterozygous females between 6-12 weeks old except Nr4a1^RFP^, Cd74^tdTomato^, Tmem119^GFP/GFP^, and Timd4^Cre/+^Rosa^YFP^ mice. For CreERT2 induction in adult mice, 50 mg/kg tamoxifen (TAM) (Millipore Sigma, T5648-5G) in corn oil was administered intraperitoneally on day -2 and day -1, and tissues were collected and processed on day 0. All procedures were approved by the Institutional Animal Care and Use committee at Northwestern University. Schematic of modified fate-mapping mouse models (created in BioRender.com).

### Tissue preparation and flow cytometric analysis

Single-cell suspensions of PB were prepared as previously described ^6,8^. 90 µL of PB, obtained via submandibular or cardiac puncture collection, was incubated with Fc Block at 4°C for 20 minutes, and then stained with an antibody cocktail for 30 minutes at 4°C (R718 CD45 (BD Biosciences, 567075), BV421 CD45.2 (BD Biosciences, 109831), BV711 Ly6G (BD Biosciences, 563979), PE-CF594 SiglecF (BD Biosciences, 562757), BV605 SiglecF (BD Biosciences, 740388), BB700 CD11b (BD Biosciences, 566416), APC CD4 (BD Biosciences, 553051), APC CD8a (BD Biosciences, 553035), APC CD19 (BD Biosciences, 550992), APC NK1.1 (BD Biosciences, 550627), PE CD115 (Invitrogen, 12-1152-82), APC Cy7 (BD Biosciences, 560596), PE Cy7 CD62L (BioLegend, 104418)). Red blood cells were lysed at room temperature for 10 minutes using FACS Lyse (BD Biosciences, 349202) diluted 1:1 in sterile water. Optimized protocol of intravascular injection (I.V.) labeling based on previous study to differentiate intra- and extra-vascular monocytes was used ^8^. Conjugated BUV661 anti-CD45 (BD Biosciences, 612975) antibody was administered to mouse via I.V. administration at 0.2 mg/mL in 200 μL DPBS after mice were lightly anesthetized with controlled isoflurane inhalation (Covetrus, 029405). Mice were then monitored for 4 minutes under housing environment before euthanasia via CO_2_ inhalation. Treml4 was used as a proxy for circulating monocytes instead of I.V. CD45 in several mouse lines as previously described ^8^. All joints from mice were dissected by cutting above the synovium and below the tibia area to isolate synovial tissues while minimizing contamination from surrounding connective tissue and processed as previously described ^6,8^. Skins and toes were removed from the dissected ankles. Each dissected ankle was flushed with 2.5 mL HBSS with Ca^2+^ and Mg^2+^ (Gibco, 14025-092, 5 mL per mouse) to get rid of the exposed bone marrow. Ankle digestion buffer was made with 2.4 mg/mL dispase II (Roche, 4942078001), 2 mg/mL collagenase D (Roche, 11088866001), 0.2 mg/mL DNAse I (Roche, 10104159001) in HBSS with the final pH adjusted to 7.2 – 7.4 for improved cell recovery. Synovial tissue was infused with 1.5 mL digestion buffer per ankle. Ankles were then incubated at 37°C for 1 hour with consistent shaking at 200 rpm. Digested ankles were then gently pressed against 40 μm nylon cell strainers (FALCON, 352340) to harvest as many synovial single-cells as possible without breaking the ankles. Two hundred microliters of BD Pharm Lyse lysing buffer (BD Biosciences, 555899) was used to lyse red blood cells for 1 minute. eFluor^TM^ 506 Fixable Viability Dye (Invitrogen, 65-0886-14) was used to stain dead synovial cells for 15 minutes. Unspecific antibody binding was avoided by incubating the single cell suspension using CD16/32 Fc Block (BD Biosciences, 553142) for 20 minutes at 4°C. Synovial cells were then stained with antibodies (R718 CD45 (BD Biosciences, 567075), BB700 CD11b (BD Biosciences, 566416), BV711 Ly6G (BD Biosciences, 563979), PE-CF594 SiglecF (BD Biosciences, 562757), BV605 SiglecF (BD Biosciences, 740388), FITC SiglecF (BioLegend, 562757), BV786 CD64 (BD Biosciences, 741024), PE Cy7 MHCII (BioLegend, 107630), APC Cy7 (BD Biosciences, 560596), AF647 CD177 (BD Biosciences, 566599), BUV737 Treml4 (BD Biosciences, 755275) for 30 minutes at 4°C. To preserve cell integrity, samples were incubated in a 2% paraformaldehyde solution (Electron Microscopy Sciences, 15713-S) in PBS at room temperature for 15 minutes, shielded from light. Analytical single-cell suspension was acquired on BD FACSymphony A5.1 Analyzer (BD Biosciences) or BD FACSymphony A5.2 Spectral Analyzer (BD Biosciences) at RHLCCC Flow Core, Northwestern University. Fluorescence minus one (FMO) controls were utilized as needed to define gating boundaries. FlowJo V10 was used for compensation and analysis of flow-cytometry data with spectral configuration and unmixing if needed.

### Single-cell (SC) RNA sequencing

For the single-cell analysis, cellular indexing of transcriptomes and epitopes by sequencing (CITE-seq) data of sorted CD45^+^CD11b^+^Ly6G^-^SiglecF^-^CD64^+^ (SC RNA: GSM7056605, ADT: GSM7056606) and CD45^+^CD11b^+^Ly6G^-^SiglecF^-^CD64^-^MHCII^-^ (SC RNA: GSM7056601, ADT: GSM7056602) was acquired from Montgomery, *et al.*, ^8^. FASTQs were re-processed in house to take advantage of the latest CellRanger pipeline. The mm10 (mm10-2020-A) mouse reference genome was used for reading processing and alignment using mkfastq and count commands of Cell Ranger (v 7.1.0). Pre-processing and analysis were performed in Seurat (v 5.1.0) in R (v 4.2.3) (see table below). Low quality cells with less than 6000 UMIs, or more than 7.5% mitochondrial genes were removed. scDblFinder (v 1.12.0) was used to call and remove doublets with an expected doublet formation rate of 5%. The RNA assay was transformed using Seurat’s log-normalization method. The ADT assay was normalized using the centered log ratio method with margin = 2 to normalize within each cell. Cell cycle scores were calculated using mouse orthologs of Seurat-provided S and G2M phase gene lists obtained through biomaRt (v 2.54.1, Dec. 2021 human and mouse archives).

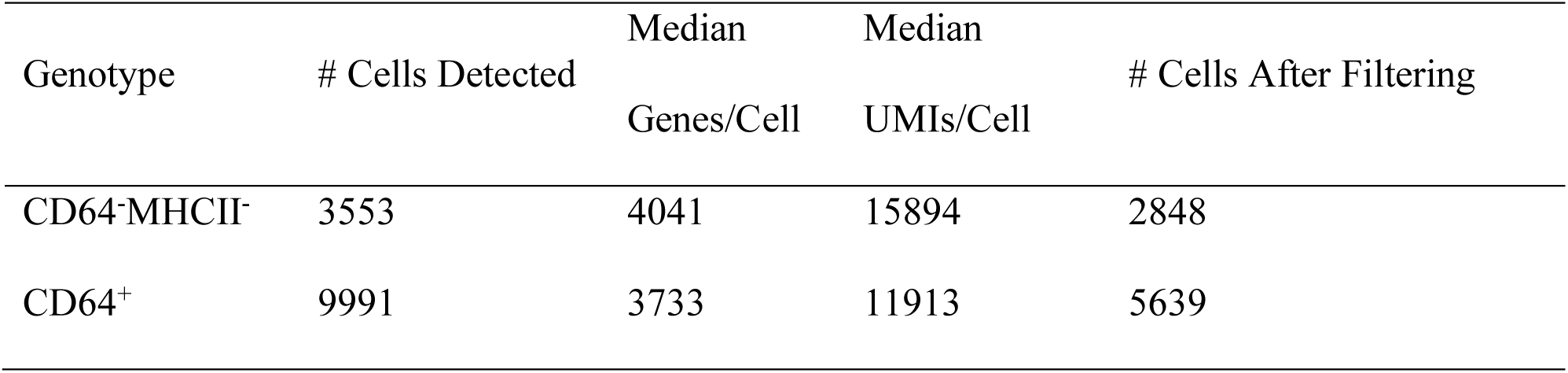

For the purposes of dimensionality reduction and clustering, SCTransform normalization was carried out on the RNA assay using 750 variable genes and regressing out S and G2M phase module scores. The joint UMAP of CD64^+^ and CD64^-^MHCII^-^ was constructed using RunUMAP with top 10 PCs. For sub-clustering of the CD64^-^MHCII^-^ dataset, FindNeighbors (14 PCs; 20 k-nearest neighbors) and FindClusters (resolution=0.1) were used. For the CD64^+^ dataset, FindNeighbors (16 PCs; 10 k-nearest neighbors) and FindClusters (resolution=0.25) were used. De-novo clusters were annotated based on canonical genes, ADT expression, and marker identified by the FindAllMarkers function with an adjusted p-value threshold of <0.05, determined by the Wilcoxon test, and adjusted using the Benjamini-Hochberg method, with all other parameters set to their default values. Plots of gene expression and reporter levels for fate-mapping markers were visualized in a web application using RShiny (v 1.9.1).

### Quantification and statistical analysis

All fate-mapping analyses were performed using GraphPad Prism (v 10.3.0). Data are presented as means ± standard deviation (SD). A total of three blood samples (from two mouse models) and five ankle samples (from five mouse models) were excluded because they contained at least one datapoint outside the two SD range in any detected population.

## RESULTS

### Gating strategy to identify PB and synovial monocyte and macrophage populations

We and others have shown that peripheral blood (PB) and synovial myeloid populations are heterogeneous in mice and humans ^1,6,8,35,36^. To complement these studies, we devised a gating strategy to analyze monocytes and macrophages using multiparameter flow cytometry on single cell suspensions from PB and digested synovial tissue. When processing blood, we first removed counting beads and doublets, then focused on CD45^+^CD11b^+^Ly6G^-^SiglecF^-^CD115^+^CD4^-^CD8^-^ CD19^-^NK1.1^-^. We identified classical monocytes (CM, Ly6C^+^CD62L^+^), intermediate monocytes (IM, Ly6C^int^CD62L^-^), and non-classical monocytes (NCM, Ly6C^-^CD62L^-^) in PB (Supplemental Fig 1).

For synovial tissue, the identification of myeloid cells is analogous to the gating strategy in PB and similar to our recent study that identified tissue resident monocytic cells (TRMC) ^8^. After excluding counting beads, doublets and dead cells, we focused on the CD45^+^ (hematopoietic cells), CD11b^+^ myeloid cells and then removed neutrophils (Ly6G^+^), eosinophils (SiglecF^+^), and dendritic cells (CD64^-^MHCII^+^) (Fig 1A). NK cells in the synovial tissue represent less than 1% of the CD45^+^ and are CD11b^neg-low^ (Fig 1B). Macrophages were identified by CD64^+^, while TRMC were I.V.CD45^-^Treml4^-^Ly6C^-^CD177^-^, CM were Ly6C^+^CD177^+/-^, and NCM were I.V.CD45^+^ Treml4^-^Ly6C^-^CD177^-^ with CD177 used to exclude Ly6G negative neutrophils ^37^ (Fig 1A, C)

**Figure 1.**
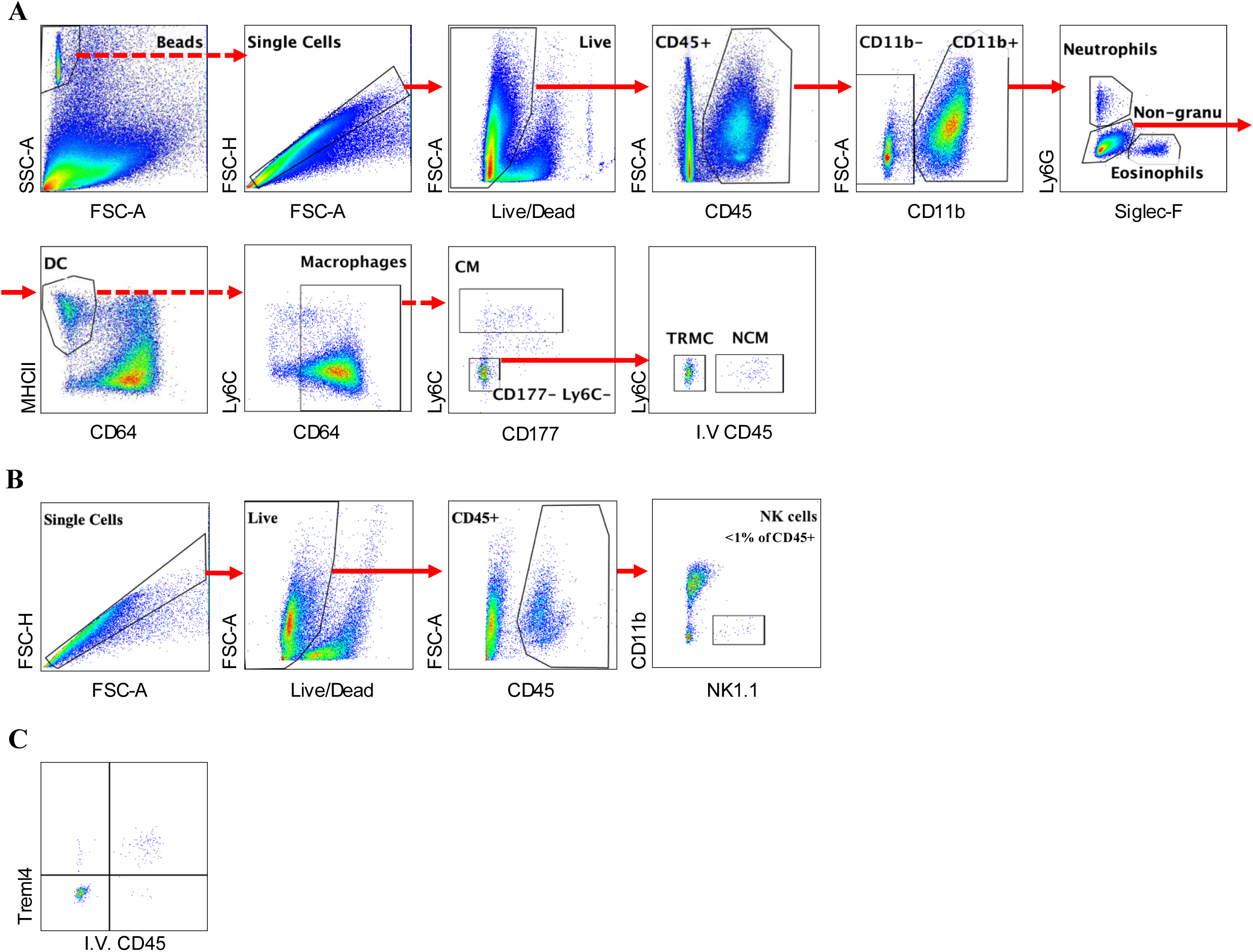
Flow cytometry gating strategy of cells in mouse synovial tissue. (A) Mouse synovial tissue gating strategy for the identification of eosinophils, neutrophils, dendritic cells (DCs), macrophages, tissue-resident monocytes-lineage cells (TRMCs), non-classical monocytes (NCMs), and classical monocytes (CMs). (B) Natural killer (NK) cells in synovial tissue are CD11b^-^. (C) Comparable level of staining with I.V. anti-CD45 antibody and anti-Treml4 antibody on NCM in synovial tissue. Solid arrows indicate positive gating, while dashed arrows indicate negative gating.

### Overview of 18 different myeloid associated fate-mapping and reporter mouse models

To better understand the efficacy and specificity of different myeloid fate-mapping and reporter mouse models in murine synovial tissue, we accumulated 18 murine strains that are considered specific for myeloid cells (Fig 2 and Table 1). We then measured the tracing ability of these myeloid promoters in PB and synovial tissue. CX3CR1 is a widely used marker to differentiate CM (CX3CR^low-neg^) and NCM (CX3CR1^+^) and is expressed in subpopulations of macrophages ^5,6,38^. Moreover, the Cx3cr1 promoter has been shown to be active in embryonic macrophage precursors ^5,38,39^. We compared YFP and GFP expressions in PB and in the synovium using Cx3cr1^CreERT2/+^Ai3^YFP^, Cx3cr1^Cre/+^Ai3^YFP^, and Cx3cr1^GFP/+^ knock-in mice. TAM was injected each day for two days prior to euthanasia for Cx3cr1^CreERT2/+^Ai3^YFP^ mice. As expected, all monocyte subsets (CM, IM, and NCM) displayed high levels of GFP (>90%) in PB from Cx3cr1^GFP/+^ mice (Supplemental Fig 2A). There was minimal GFP detected in PB lymphocytes, neutrophils and eosinophils from Cx3cr1^GFP/+^ mice (Supplemental Fig 2A). These data are similar to Cx3cr1^Cre/+^Ai3^YFP^ mice except we detected reduced levels of expression in YFP from lymphocytes (13%), neutrophils (8%) and eosinophils (6%) (Supplemental Fig 2B). In contrast, only a fraction of the PB CM (21%) were YFP^+^, while the majority of NCM and IM (>90%) were YFP^+^ in PB from TAM-treated Cx3cr1^CreERT2/+^Ai3^YFP^ mice (Supplemental Fig 2C). A low but not negligible expression of YFP was detected in CD45^+^ cells (0.27%) from PB in corn oil-treated Cx3cr1^CreERT2/+^Ai3^YFP^ mice ^40^ (Supplemental Fig 4A). Together these are consistent with previous reports ^36,38^.

**Figure 2.**
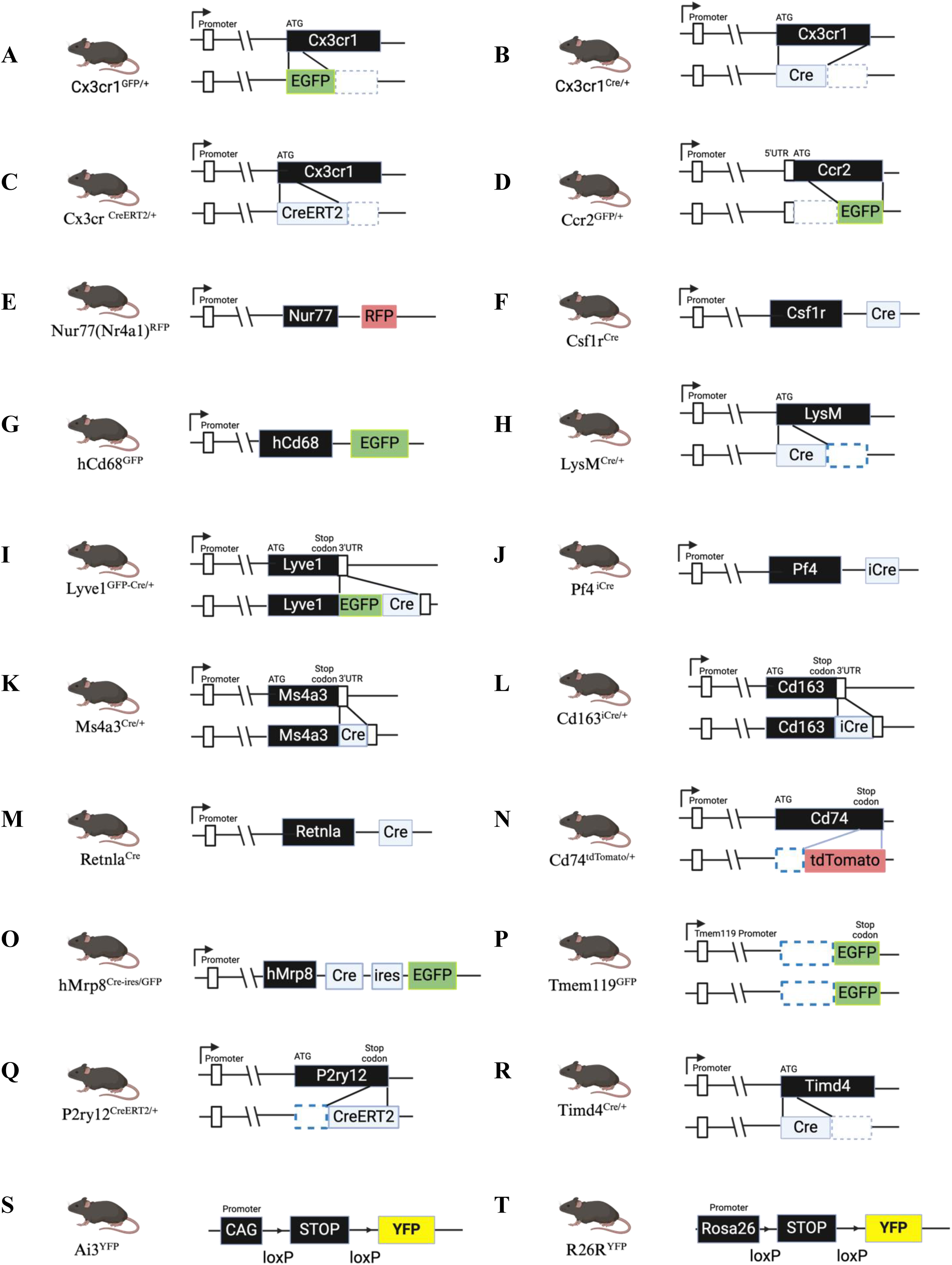
Schematic of modified fate-mapping mouse models. The schematic of (A) Cx3cr1^CreERT2/+^, (B) Cx3cr1^Cre/+^, (C) Cx3cr1^GFP/+^, (D) Ccr2^GFP+^, (E) Nr4a1^RFP^, (F) Csf1r^Cre^, (G) hCD68^GFP^, (H) LysM^Cre/+^, (I) Lyve1^GFP-Cre/+^, (J) Pf4^iCre^, (K) Ms4a3^Cre/+^, (L) Cd163^iCre/+^,(M) Retnla^iCre^,(N) Cd74td^Tomato/+^, (O) hMrp8^Cre-ires/GFP^, (P) Tmem119^GFP/GFP^, (Q) P2ry12^CreERT2/+^, (R) Timd4^Cre/+^, (S) Ai3^YFP^, and (T) R26^YFP^ mouse model.

**Table 1.** Insertion types and references of the fate mapping mouse lines.

| **Genotypes** | **Insertion** | **References** | **Jax #** |
| --- | --- | --- | --- |
| Cx3cr1<sup>GFP/+</sup> | Knock-in | Yona S *et al.*, 2013 | 005583 |
| Cx3cr1<sup>Cre/+</sup> | Knock-in | Yona S *et al.*, 2013 | 025524 |
| Cx3cr1<sup>CreERT2/+</sup> | Knock-in | Yona S *et al.*, 2013 | 020940 |
| Ccr2<sup>GFP/+</sup> | Knock-in | Satpathy AT , *et al.*, 2013 | 027619 |
| Nur77(Nr4a1)<sup>RFP</sup> | Transgenic | Hemmers S , *et al.*, 2019 | 034918 |
| Csf1r<sup>Cre</sup> | Transgenic | Loschko J , *et al.*, 2016 | 029206 |
| hCD68<sup>GFP</sup> | Transgenic | Iqbal AJ *et al.*, 2014 | 026827 |
| LysM<sup>Cre/+</sup> | Knock-in | Clausen BE , *et al.*, 1999 | 004781 |
| Lyve1<sup>GFP-Cre/+</sup> | Knock-in | Pham *et al.*, 2010 | 012601 |
| Pf4<sup>iCre</sup> | Transgenic | Tiedt R *et al.*, 2006 | 008535 |
| Ms4a3<sup>Cre/+</sup> | Knock-in | Liu Z , *et al.*, 2019 | 036382 |
| Cd163<sup>iCre/+</sup> | Knock-in | Etzerodt A, *et al.*, 2019 | Not applicable |
| Retnla<sup>Cre</sup> | Transgenic | Krljanac B, *et al.*, 2019 | Not applicable |
| P2ry12<sup>CreERT2/+</sup> | Knock-in | Mckinsey GL, *et al.*, 2020 | 034727 |
| Tmem119<sup>GFP/GFP</sup> | Knock-in | Kaiser T, *et al.*, 2019 | 031823 |
| Timd4<sup>Cre/+</sup> Rosa<sup>YFP</sup> | Knock-in | Prise IE, *et al.*, 2023 | No applicable |
| hMRP8<sup>Cre/ires-GFP</sup> | Transgenic | Passegue E , *et al.*, 2004 | 021614 |
| Cd74<sup>tdTomato/+</sup> | Knock-in | Harrington EP, *et al.*, 2023 | Not applicable |
| Ai3 | Transgenic | Madisen L , *et al.*, 2010 | 007903 |
| R26R-YFP | Knock-in | Srinivas S, *et al.*, 2001 | 006148 |

In the synovium, GFP was highly expressed in the CM (90%) and NCM (>90%) that are retained in the vasculature from the digested ankles from Cx3cr1^GFP/+^ mice (Fig 3A). Around 50% of the dendritic cells were GFP^+^, while macrophages (26%) and TRMC (43%) displayed reduced levels of GFP in the synovium of Cx3cr1^GFP/+^ mice (Fig 3A). The CD45^+^CD11b^-^ population expressed low levels of GFP and the eosinophils as well as neutrophils were negative for GFP (Fig 3A). The synovial data from Cx3cr1^GFP/+^ mice differs greatly from the Cx3cr1^Cre/+^Ai3^YFP^ mice. We detected YFP^+^ in all synovial hematopoietic cells in Cx3cr1^Cre/+^Ai3^YFP^ mice; 58% in CD45^+^CD11b^-^, 59% in eosinophils, 64% in neutrophils, 93% in DCs, 91% in macrophages, 73% in CM, 87% in NCM, and 79% in TRMC (Fig 3B). In contrast, Cx3cr1^CreERT2/+^Ai3^YFP^ mouse profiling data were more similar to Cx3cr1^GFP/+^ mouse model, albeit markedly fewer YFP^+^ cells in DC (8%), macrophage (23%), CM (13%), NCM (80%), and TRMC (17%) (Fig 3C). We did not detect any YFP above baseline in the CD45^+^CD11b^-^, eosinophils, and neutrophils from TMX treated Cx3cr1^CreERT2/+^Ai3^YFP^ mice (Fig 3A). A notable observation is the minimal YFP expression in the synovial CD45^+^ cells from the corn oil-treated Cx3cr1^CreERT2/+^Ai3^YFP^ mice (4.5%), which is consistent with previously published results ^40–43^ (Supplemental Fig 4B).

**Figure 3.**
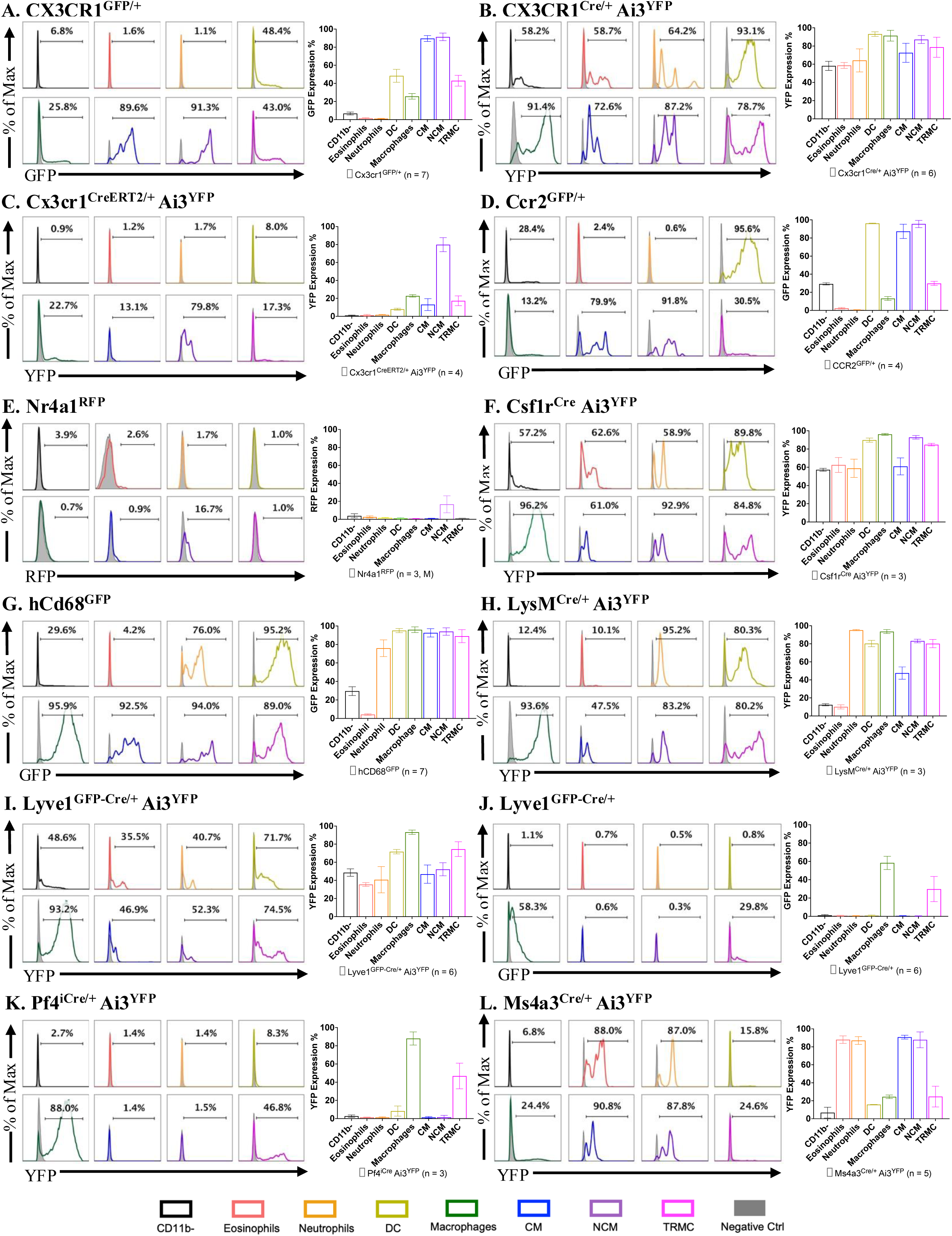
Quantification of reporter-gene positive cells in synovial immune population. Histogram of a representative mouse (percent indicates mean reporter gene-positive across all mice) and bar graph (mean ± SD) summarizing frequencies of reporter gene-positive synovial cells in twelve reporter and fate-mapping models. (A) Cx3cr1^GFP/+^ (n= 7; C57BL/6 mice were used as negative controls). (B) Cx3cr1^Cre/+^Ai3^YFP^ (n=6; C57BL/6 or Ai3^YFP^ mice were used as negative controls). (C) Cx3cr1^CreERT2/+^Ai3^YFP^ (n=4; Ai3^YFP^ mice were used as negative controls; 50 mg/kg tamoxifen (TAM) in corn oil was administered intraperitoneally on day -2 and day -1, and tissues were collected and processed on day 0). (D) Ccr2^GFP/+^ (n=4; C57BL/6 mice were used as negative controls). (E) Nr4a1^RFP^ (n=3, males; C57BL/6 mice were used as negative controls). (F) Csf1r^Cre^ Ai3^YFP^ (n=3; Ai3^YFP^ mice were used as negative controls). (G) hCD68^GFP^ (n =7; TRMC was identified using I.V. CD45 in 5 of the mice, and Treml4 in 2 of the mice; C57BL/6 mice were used as negative controls). (H) LysM^Cre/+^Ai3^YFP^ (n=3; C57BL/6 mice were used as negative controls). (J) Lyve1^GFP-Cre/+^Ai3^YFP^ (n = 6; 5 of the mice received I.V. CD45 for the identification of TRMC, in 1 mouse Treml4 was used for the identification of TRMC; Ai3^YFP^ mice were used as negative controls), (I) Lyve1^GFP-Cre/+^ (n=6; C57BL/6 mice were used as negative controls). (K) Pf4^iCre/+^Ai3^YFP^ (n=3; Ai3^YFP^ mice were used as negative controls). (L) Ms4a3^Cre/+^Ai3^YFP^ (n=5; C57BL/6 or Ai3^YFP^ mice were used as negative controls). (M) Cd163^Cre/+^Ai3^YFP^ (n=4; Ai3^YFP^ mice were used as negative controls). (N) Retnla^Cre^Ai3^YFP^ (n=2; Ai3^YFP^ mice were used as negative controls). (O) Cd74^tdTomato/+^ (n=6, 4 females, 2 males; C57BL/6 mice were used as negative controls). (P) hMrp8^Cre/ires-GFP^Ai3^YFP^ (n=5; Ai3^YFP^ mice were used as negative controls). (Q) Tmem119^GFP/GFP^ (n=4; C57BL/6 or Ai3^YFP^ mice were used as negative controls). R) Histogram and bar graph of P2ry12^CreERT2^/+ Ai3^YFP^ (n = 5; P2ry12^ERCre/+^Ai3^YFP^ mouse without treatment or C57BL/6 mice were used as negative controls; 50mg/kg TAM in corn oil was administered intraperitoneally on day -2 and day -1, and tissues were collected and processed on day 0). (S) Tim4^Cre/+^R26^YFP/+^ (n=4, males; R26^YFP^ mice were used as negative controls).

Ccr2 and Nr4a1 are known to be crucial for the development of CM and NCM, respectively ^1^. Almost all PB CM, IM, and NCM were GFP^+^ in Ccr2^GFP/+^ knock-in mice, as NCM are derived from CM (Supplemental Fig 2D). There was minimal expression of GFP in lymphocytes, neutrophils or eosinophils from CCR2^GFP/+^ mice (Supplemental Fig 2D). The Nr4a1^RFP^ transgenic mice only displayed RFP^+^ in PB IM (29%) and NCM (73%) (Supplemental Fig 2E). GFP expression was detected in the vast majority DCs (96%), CM (80%), and NCM (92%), while synovial macrophages (13%) and TRMC (31%) had markedly fewer GFP^+^ cells in the synovium of Ccr2^GFP/+^ mice (Fig 3D). Only the NCM (17%) displayed RFP positivity in the synovium of Nr4a1^RFP^ mice (Fig 3E).

One of the central issues of general myeloid specific promoters is their ubiquitous expression across multiple myeloid subtypes of cells. Csf1r, hCd68 and LysM, are commonly used promoters to express Cre or a reporter such as GFP to yield expression in monocytes and macrophages ^44–46^. Lyve1 expression was shown to denote perivascular location for tissue resident macrophages ^28,47,48^, while Pf4 reporter mice demonstrate expression in megakaryocytes as well as monocyte/macrophage lineage cells ^49^. More recently, Ms4a3 reporter mice identified monocytes and granulocytes but not lymphocytes ^30^. The Csf1r promoter was active in all circulating leukocytes such that YFP was expressed in lymphocytes (69%), eosinophils (38%), neutrophils (66%), CM (99%), IM (98%), and NCM (99%) in PB from Csf1r^Cre^Ai3^YFP^ mice (Supplemental Fig 2F). As expected, PB from hCD68^EGFP^ mice only expressed GFP in neutrophils (49%), CM (95%), IM (97%), and NCM (96%) (Supplemental Fig 2G). Similarly, LysM^Cre^Ai3^YFP^ and Lyve1^Cre^Ai3^YFP^ mice also displayed YFP^+^ in lymphocytes (14%, 35%), eosinophils (16%, 26%), neutrophils (70%, 6%), CM (78%, 63%), IM (82%, 60%), and NCM (65%, 100%) in PB, respectively (Supplemental Fig 2H-I). Surprisingly, Lyve1^Cre^ mice, which also express GFP as well as Cre from the Lyve1 promoter, had undetectable GFP in all populations from PB (Supplemental Fig 2J). Pf4^iCre^Ai3^YFP^ mice also failed to express YFP in PB leukocytes (Supplemental Fig 2K), while Ms4a3^Cre/+^Ai3^YFP^ mice showed moderate expression of YFP in eosinophils (32%) and neutrophils (24%) but higher number of positive cells in CM (84%), IM (78%), and NCM (88%) from PB (Supplemental Fig 2L).

All synovial leukocyte populations expressed YFP (>50%) in Csf1r^Cre^Ai3^YFP^ mice but the vast majority of DC (90%), macrophage (96%), NCM (93%), and TRMC (85%) were YFP^+^ (Fig 3F). hCD68^GFP^ also displayed ubiquitous expression of GFP in all leukocytes except eosinophils and had highest expression in DC (95%), macrophage (96%), CM (93%), NCM (94%), and TRMC (89%) (Fig 3G). LysM^Cre^Ai3^YFP^ mice also showed YFP^+^ in all leukocytes, although at low expression levels in lymphocytes (12%) and eosinophils (10%) (Fig 3H). Similarly, Lyve1^Cre^Ai3^YFP^ mice showed YFP^+^ in all leukocytes, with the highest expressions in DC (72%), macrophage (93%), and TRMC (72%) yielded the greatest number of positive cells (Fig 3I). GFP expression was detected only in macrophages (58%) and TRMC (30%) from Lyve1^GFP-Cre/+^ mice (Fig 3J), albeit at lower levels than YFP from Lyve1^Cre/+^Ai3^YFP^ mice. Pf4^iCre^Ai3^YFP^ mice had a more restrictive expression of YFP as 88% of synovial macrophage and 47% of TRMC were positive, while the other leukocyte populations had negligible YFP expression (Fig 3K). Similar to PB, YFP was detected in all synovial immune populations with the highest being eosinophils (88%), neutrophils (87%), CM (91%) and NCM (88%) from Ms4a3^Cre/+^Ai3^YFP^ mice (Fig 3L).

CD163 has been reported to be highly expressed in red-pulp macrophages in the spleen ^50^, while Retnla^Cre^ mouse efficiently traces white adipose tissue, peritoneal, lung, and intestinal macrophages, as well as granulocytes ^34^. Although only low level of YFP was detected in lymphocytes (31%), CM (8%), IM (13%), and NCM (13%) from PB in Cd163^iCre/+^Ai3^YFP^ mice (Supplemental Fig 3M), synovial DC (90%), macrophages (92%), and TRMC (80%) expressed high levels of YFP, with 37% of lymphocytes displaying YFP^+^ (Fig 4M). None of the PB cells expressed YFP in Retnla^Cre^Ai3^YFP^ mice (Supplemental Fig 3N), consistent with published results^34^. Moderate level of YFP^+^ cells were detected in synovial eosinophils (34%), DC (51%), macrophages (47%), and TRMC (25%) in Retnla^Cre^Ai3^YFP^ mice (Fig 4N).

**Figure 4.**
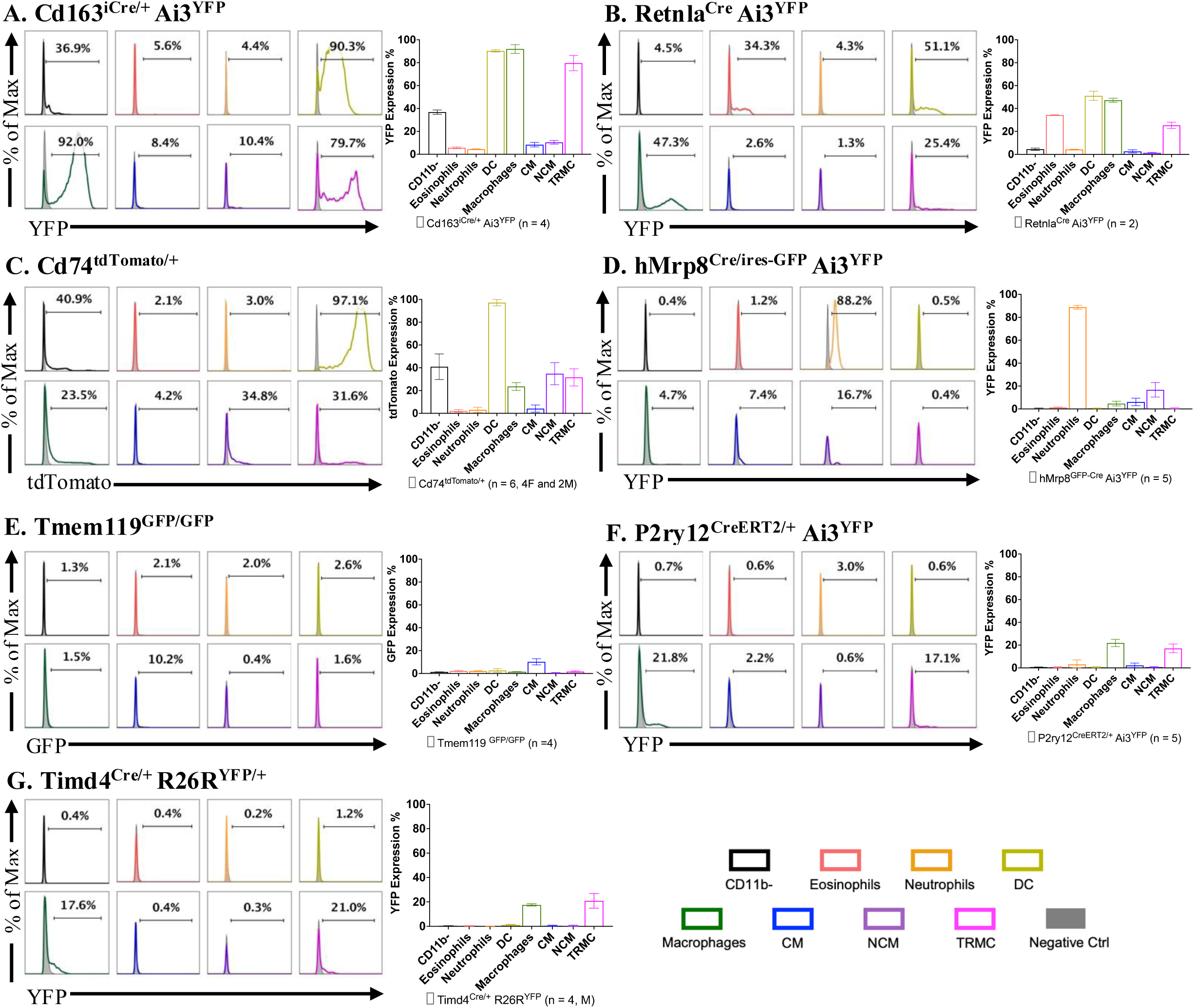
Quantification of reporter-gene positive cells in synovial immune population. Histogram of a representative mouse (percent indicates mean reporter gene-positive across all mice) and bar graph (mean ± SD) summarizing frequencies of reporter gene-positive synovial cells in seven reporter and fate-mapping models. (A) Cd163^Cre/+^Ai3^YFP^ (n=4; Ai3^YFP^ mice were used as negative controls). (B) Retnla^Cre^Ai3^YFP^ (n=2; Ai3^YFP^ mice were used as negative controls). (C) Cd74^tdTomato/+^ (n=6, 4 females, 2 males; C57BL/6 mice were used as negative controls). (D) hMrp8^Cre/ires-GFP^Ai3^YFP^ (n=5; Ai3^YFP^ mice were used as negative controls). (E) Tmem119^GFP/GFP^ (n=4; C57BL/6 or Ai3^YFP^ mice were used as negative controls). F) Histogram and bar graph of P2ry12^CreERT2^/+ Ai3^YFP^ (n = 5; P2ry12^ERCre/+^Ai3^YFP^ mouse without treatment or C57BL/6 mice were used as negative controls; 50mg/kg TAM in corn oil was administered intraperitoneally on day -2 and day -1, and tissues were collected and processed on day 0). (G) Tim4^Cre/+^R26^YFP/+^ (n=4, males; R26^YFP^ mice were used as negative controls).

CD74 is required for MHCII function ^31^. Almost all populations including lymphocytes (52%), CM (21%), IM (66%), and NCM (78%) in PB except for neutrophils expressed tdTomato using Cd74^tdTomato/+^ mice (Supplemental Fig 3O). However, tdTomato was almost exclusively expressed in synovial DC (94%) (Fig 4O), although synovial macrophages (24%), NCM (35%), and TRMC (32%) had lower percents of tdTomato^+^ cells (Fig 4O). While MRP8 is known to be expressed in macrophages ^51^, low level of YFP expression was detected in IM (22%) and NCM (18%) in PB from MRP8^Cre-ires-GFP^Ai3^YFP^ mice (Supplemental Fig 3P). In synovial tissue, MRP8^Cre-ires-GFP^Ai3^YFP^ mice displayed YFP^+^ in the neutrophil population (88%), with a small percent of positivity in NCM (17%) (Fig 4P).

Tmem119^GFP/GFP^, P2ry12^CreERT2/+^Ai3^YFP^ Timd4^Cre/+^Rosa^YFP^ mice were recently generated as macrophage reporter mice ^24,25,32^. Tmem119^GFP/GFP^ mice only expressed low level of GFP in CM and IM (<15%) from PB and failed to express GFP in any of the synovial immune populations except CM (10.2%) (Supplemental Fig 3Q, Fig 4Q). P2ry12^CreERT2^Ai3^YFP^ mice treated with TMX did not express any YFP in PB cells but exhibited YFP in synovial macrophages (22%) and TRMC (17%) (Supplemental Fig 3R, Fig 4R). A very minimal YFP expression was observed in the synovial CD45^+^ cells from the corn oil-treated P2ry12^CreERT2/+^Ai3^YFP^ mice (0.6%), which is consistent with previously published results (Supplemental Fig 4C) ^25,41,43^. Timd4^Cre^Rosa^YFP^ mice only displayed YFP^+^ in a minority of macrophages (18%) and TRMC (21%); no other leukocytes exhibited any YFP^+^ in the synovium (Fig 4S).

### Comparison between gene expression and fate-mapping in synovial myeloid subpopulations

To investigate expression of fate-mapping genes in transcriptionally defined subpopulations compared with flow analysis, we used our previously published CITE-seq data on synovial myeloid populations split into CD45^+^CD11b^+^Ly6G^-^SiglecF^-^CD64^+^ macrophages and CD45^+^CD11b^+^Ly6G^-^SiglecF^-^CD64^-^MHCII^-^ myeloid cells (Fig 5A-B) ^8^. From the latter, we identified TRMCs, a long-lived tissue-resident myeloid populations that we previously described ^8^, and six other cell populations that are distinct from macrophages including CM, NCM, cycling monocytes, TRMC, CD177^+^ neutrophil and cycling CD177^+^ neutrophil population (Fig 5A-B, Supplemental Fig 5A-D; Table S1A-B). As seen previously, CD177^+^ neutrophils express neutrophil genes but lack Ly6G expression and are consistent with the population gated out in our flow analysis (Fig 1A; Supplemental Fig 5D). Overall, expression measured by transcriptional profiling agreed with the fate-mapping results. Lyve1, Pf4, Cd163, Retnla, P2ry12, and Timd4 were expressed in TRMC and CD64^+^ cells (macrophages) but not in CM or NCM (Fig 5C). Ms4a3 was not expressed in CM, NCM nor TRMC, although the YFP was widespread in the Ms4a3^Cre^ model (Fig 5C-D). These results are consistent with the expression of Ms4a3 in progenitor cells but not maintained in terminal cells ^30^. All other genes such as Cx3cr1, Ccr2, Nr4a1, Csf1r, Lyz2, and Cd74 were detected at some level in macrophages, monocytes, and TRMC, which corresponded to the results from reporter mice in CM, NCM and TRMC (Fig 5C-D). The gene expression of Cd68 and the corresponding GFP signal in the human counterpart hCD68^GFP^ model were prevalent in macrophages and all monocyte-lineage cells (Supplemental Fig 5E-F). S100a8 (Mrp8) gene expression was absent in macrophages but widely present in CM, NCM and TRMC, while YFP expression in human counterpart hMrp8^GFP-Cre^Ai3^YFP^ mice was minimal to absent across these populations. (Supplemental Fig 5E-F).

**Figure 5.**
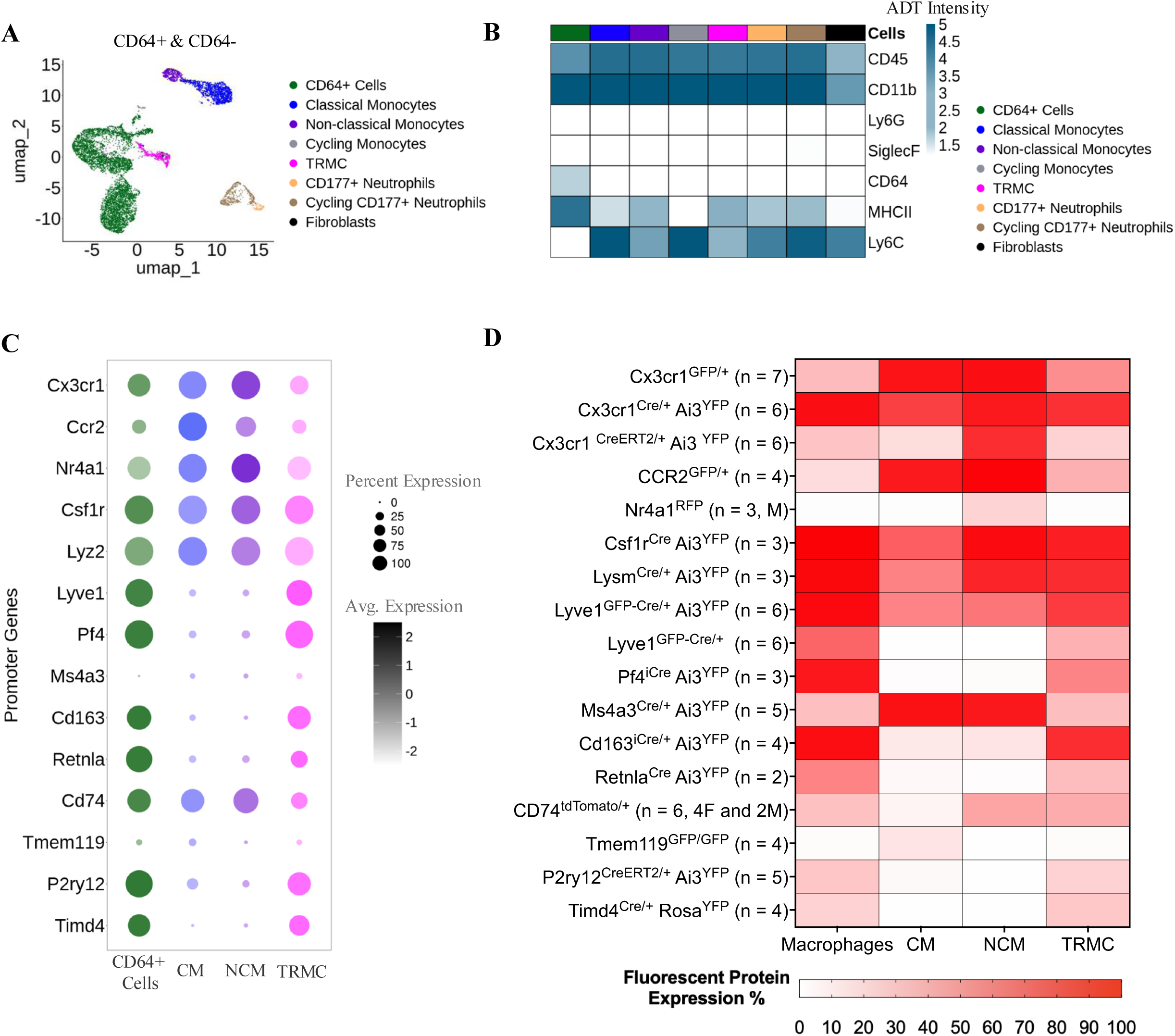
Comparison of gene expression and fate-mapping reporters in monocyte lineage subsets in the synovium. (A) Uniform manifold approximation and projection (UMAP) depicting eight cell populations from C57BL/6 CD45^+^CD11b^+^CD4^-^CD8^-^CD19^-^NK1.1^-^Ly6G^-^SiglecF^-^ CD64^+^ and CD45^+^CD11b^+^CD4^-^CD8^-^CD19^-^NK1.1^-^Ly6G^-^SiglecF^-^CD64^-^MHCII^-^ cells. (B) Heatmap of the average expression of the ADT markers of the eight cell populations identified in A. (C) Bubble plots of the average scaled gene expression of promoter genes in the CD64^+^ cells, classical monocytes (CM), non-classical monocytes (NCM), and TRMC defined by CITE-seq in A. (D) Heatmap of the percent of cells with detectable fluorescent protein in each fate-mapping model across macrophages, CM, NCM, and TRMC defined by flow cytometry.

In our reanalysis of CITE-seq data from sorted CD45^+^CD11b^+^Ly6G^-^SiglecF^-^CD64^+^ synovial macrophages ^8^, we defined six clusters which were then annotated based on the ADT levels of Cx3cr1 (CX3CR1) and H2-Aa (MHCII) protein; CX3CR1^+^MHCII^-^, CX3CR1^+^MHCII^+^, CX3CR1^-^ MHCII^-^ A, CX3CR1^-^MHCII^-^ B, CX3CR1^-^MHCII^+^ A, CX3CR1^-^MHCII^+^ B. (Fig 6A-C; Supplemental Figure 6A-B; Table S2A-B). Corresponding macrophage subsets were identified using flow cytometry with surface marker CX3CR1 and MHCII (Fig 6D-F).

**Figure 6.**
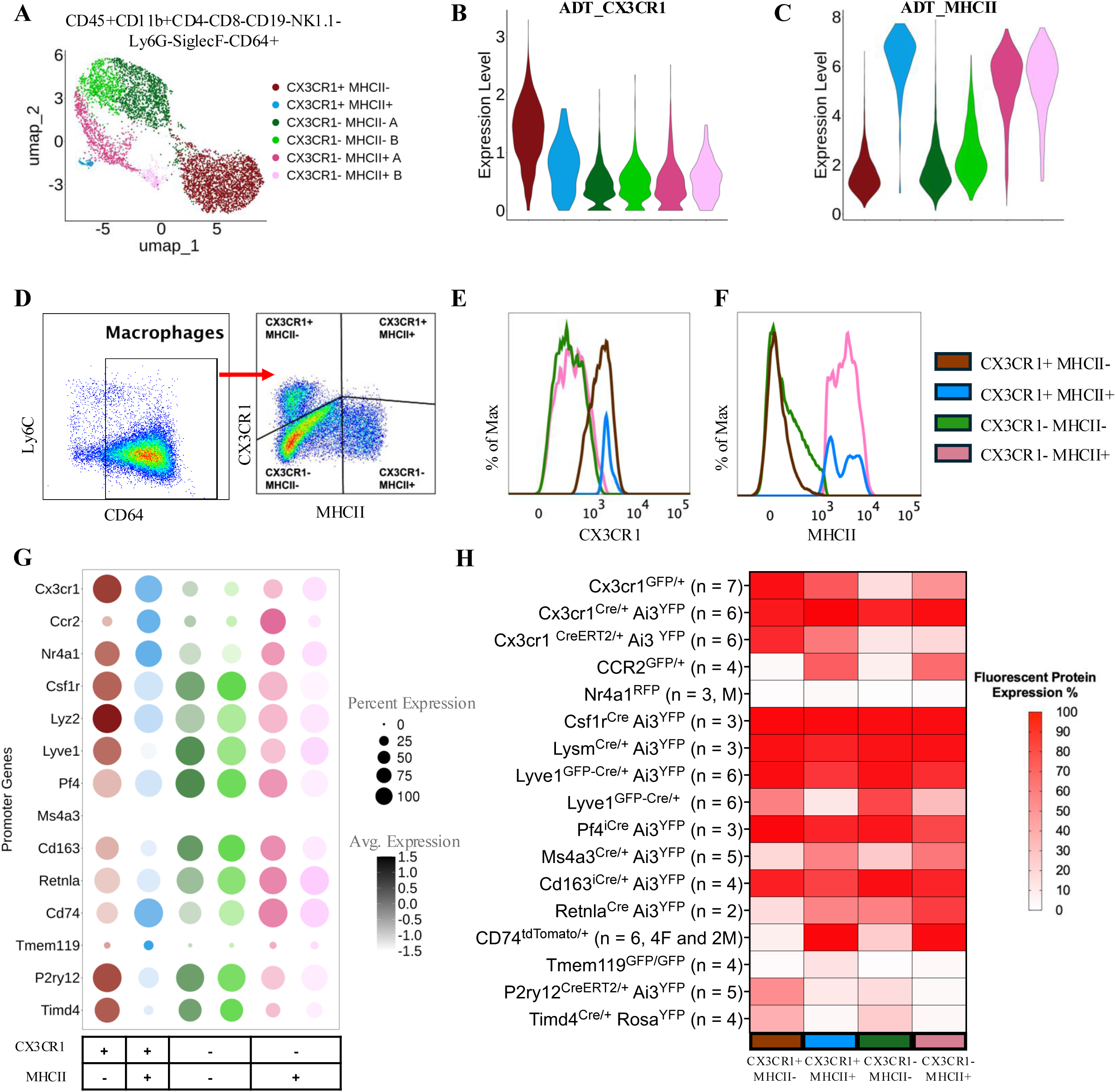
Comparison of gene expression and fate-mapping reporters in macrophage subsets in the synovium. (A) UMAP depicting six macrophage subsets of C57BL/6 CD45^+^CD11b^+^CD4^-^ CD8^-^CD19^-^NK1.1^-^Ly6G^-^SiglecF^-^CD64^+^ cells. (B) Violin plots of the ADT expression of CX3CR1 and (C) I-A-I-E across macrophage subsets. (D) Four synovial macrophages identified by flow cytometry using CX3CR1 vs. MHCII. (E) CX3CR1 surface protein expression in synovial macrophages. (F) MHCII surface protein expression in synovial macrophages. (G) Bubble plots of the average scaled gene expression of promoter genes in the six macrophage subsets defined by CITE-seq in A. (H) Heatmap of the percent of cells with detectable fluorescent protein in each fate-mapping model across the four macrophage subsets defined by flow cytometry in D.

We then compared endogenous gene expression of Cx3cr1, Ccr2, Nr4a1, Csf1r, LysM, Lyve1, Pf4, Cd163, Retnla, P2ry12, Tmem119, Timd4, and Cd74 to reporter labelling using the myeloid Cre fate mapping mice in macrophage subsets (Figure 6G-H). Cx3cr1 gene is highly expressed in the two CX3CR1^+^ subsets. However, the CX3CR1^+^MHCII^-^ population displayed the highest percent of YFP/GFP in Cx3cr^CreERT2/+^Ai3^YFP^ and Cx3cr1^GFP/+^ mice, while all subsets exhibited almost 100% of YFP^+^ in Cx3cr1^Cre/+^Ai3^YFP^ mice. CX3CR1^+^MHCII^+^ and CX3CR1^-^MHCII^+^ A populations expressed the highest levels of Ccr2, which is consistent with GFP being exclusively observed in all MHCII^+^ synovial macrophages by flow cytometry. Nr4a1 was detected in all synovial macrophage subsets yet there was minimal expression of RFP in all subsets. Csf1r and LysM genes were detected in synovial macrophage cells. These data are consistent with observation of YFP^+^ in all macrophage subsets from Csf1r^Cre/+^Ai3^YFP^ and LysM^Cre/+^Ai3^YFP^ mice. Lyve1 demonstrated a gradient of expression with the MHCII^+^ macrophage populations at the lower end. While the floxed YFP⁺ cells were detected across all CX3CR1⁺ subsets, GFP levels were markedly lower in the MHCII⁺ populations from Lyve1^GFP-Cre/+^ mice. Pf4, Cd163, and Retnla were expressed with similar patterns across macrophage subsets and YFP was ubiquitous in the Pf4 and Cd163 fate-mapping models. However, CX3CR1^+^ synovial lining macrophages displayed notably low levels of expression in Retnla^Cre^Ai3^YFP^ mice. Cd74 gene was exclusively expressed in MHCII^+^ macrophage subsets, which also exhibited high tdTomato expression from Cd74^tdTomato/+^ mice. While YFP was expressed in MHCII^+^ subsets from Ms4a3^Cre/+^Ai3^YFP^ mice, gene expression of Ms4a3 was undetectable as expected. Trem119 was minimally detected in synovial macrophages subsets at the gene expression and reporter level. P2ry12 and Timd4 were expressed primarily in MHCII^-^ cells and exhibit moderate YFP signal in the same subsets from P2ry12^CreERT2/+^Ai3^YFP^ and Timd4^Cre/+^Rosa^YFP^ mice. Cd68 and S100a8 gene expression were ubiquitously present and absent across macrophages, respectively, which matched the observed GFP/YFP levels from their human counterparts in hCD68^GFP^ and hMrp8^GFP-Cre^Ai3^YFP^ mice (Supplemental Fig 6C-D). Gene expression and reporter levels for all fate-mapping markers are available for visualization at: https://winterlab.shinyapps.io/fatemapping_macrophages/.

Since none of the known myeloid reporter mice were able to robustly distinguish individual subpopulations of TRMC or synovial macrophages, we used the CITE-seq data to search for other potential markers of myeloid subpopulations, which are more highly expressed in the target populations while showing minimal expression in the rest of the cells (Table S1A) ^8^. We identified four genes, Anxa8, Lrrc42, Nbl1, and Cuedc1 that were specifically expressed only in TRMC but not in total synovial macrophages, monocytes or neutrophils (Fig 7A) using the CD45^+^CD11b^+^Ly6G^-^SiglecF^-^CD64^-^MHCII^-^ dataset. However, following subclustering of synovial macrophages, only Lrrc42, Nbl1 and Cuedc1 were uniquely expressed in TRMC as Anxa8 was lowly expressed in CX3CR1^-^MHCII^+^ B macrophage subset, which was a minor population, when compared with the macrophage subsets from the CD45^+^CD11b^+^Ly6G^-^SiglecF^-^ CD64^+^ population (Fig 7A). Among the individual macrophage subpopulations, Ecscr/Chst1 (Fig 7B), Tnxb/Ildr2 (Fig 7D), Enpep/Chp2 (Fig 7E), Ndnf/Lsr (Fig 7F) and Arg1/Flt1 (Fig 7G) were exclusively expressed in CX3CR1⁺MHCII⁻, CX3CR1⁻MHCII⁻ A, CX3CR1⁻MHCII⁻ B, CX3CR1⁻MHCII⁺ A, and CX3CR1⁻MHCII⁺ B macrophage subsets, respectively. While Plbd1 and Sema4d were detected in CX3CR1⁺MHCII⁺ macrophages, monocytes and CD177^+^ neutrophils also expressed these genes (Fig 7C). Taken together, these data demonstrate that current myeloid reporter mice have the potential to target macrophages depending on the tissue but may also impact other immune cells. Moreover, by taking advantage of known single-cell or bulk datasets as well as curated atlases, we can identify potential promoters that are uniquely expressed in individual cell subtypes, which will greatly improve the generation of new reporter mice.

**Figure 7.**
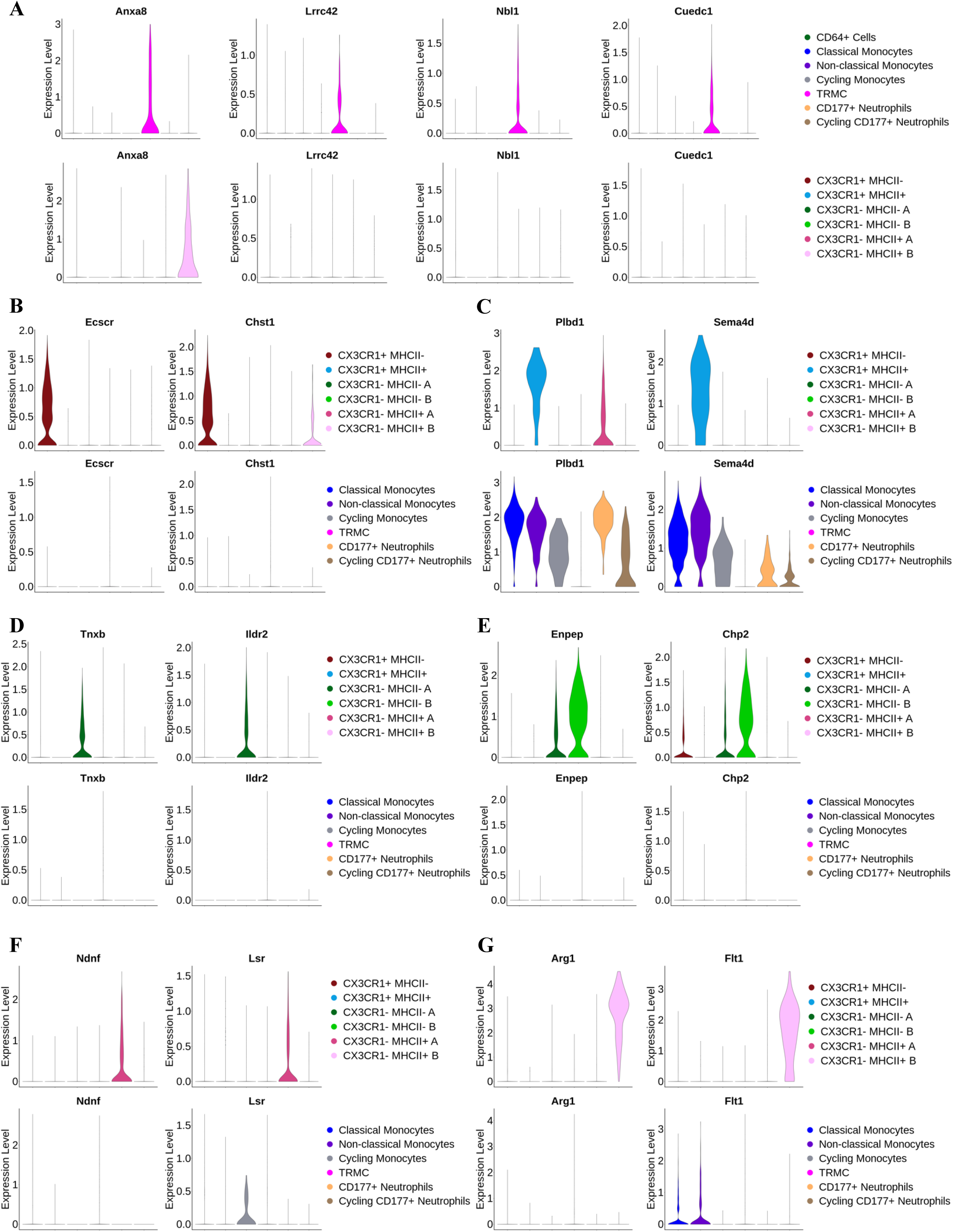
Expression of candidate gene expression markers in TRMC and synovial macrophage subsets. (A) Violin plots for the expression of potential TRMC markers in the seven myeloid populations in the merged CD45^+^CD11b^+^CD4^-^CD8^-^CD19^-^NK1.1^-^Ly6G^-^SiglecF^-^CD64^-^ MHCII^-^ (CD64^-^) and CD45^+^CD11b^+^CD4^-^CD8^-^CD19^-^NK1.1^-^Ly6G^-^SiglecF^-^CD64^+^ (CD64^+^) cells and in the six macrophage subsets from the CD64^+^ cells. (B) Violin plots for the expression of potential markers for CX3CR1^+^MHCII^-^ macrophage subset in the CD64^+^ cells and in the CD64^-^ cells. (C) Violin plots for the expression of potential markers for CX3CR1^+^MHCII^+^ macrophage subset in the CD64^+^ cells and in the CD64^-^ cells. (D) Violin plots for the expression of potential markers for CX3CR1^-^MHCII^-^ A macrophage subset in the CD64^+^ cells and in the CD64^-^ cells. (E) Violin plots for the expression of potential markers for CX3CR1^-^MHCII^-^ B macrophage subset in the CD64^+^ cells and in the CD64^-^ cells. (F) Violin plots for the expression of potential markers for CX3CR1^-^MHCII^+^ A macrophage subset in the CD64^+^ cells and in the CD64^-^ cells. (G) Violin plots for the expression of potential markers for CX3CR1^-^MHCII^+^ B macrophage subset in the CD64^+^ cells and in the CD64^-^ cells.

## DISCUSSION

Our goal in this study is to provide a comparative analysis of known reporter mice for fate-mapping of murine myeloid cells with respect to the development of arthritis. Previous studies examined the fidelity of 4 myeloid reporter mice in lung ^45^ and brain ^41,43^ as well as 13 reporter mice in blood, spleen, peritoneum and bronchioalveolar lavage ^44^. Here, we examined 18 fate-mapping and reporter mouse models, which have been previously documented to be specific for myeloid cells. To our knowledge, this is the most comprehensive study using the largest known number of reporter mice. We show that only Pf4, Tim4, P2ry12 could direct reporter expression specifically in the synovial macrophage populations but not in circulating monocytes. In contrast, Cx3cr1, Csf1r, hCD68, LysM, Lyve1, Ms4a3, Cd163, Retnla, and Cd74 reporter mice exhibit promiscuous expression in other synovial immune cells including lymphocytes, neutrophils, eosinophil and dendritic cells. To overcome overlapping expression patterns of monocyte and macrophage genes, we utilized CITE-seq on single cell suspensions from murine synovium for the identification of new macrophage specific targets. We believe that by integrating single cell analysis with the current studies thereby generating a cell atlas will bring about new and more specific targeted reporter mice that address the heterogeneity, which exists in macrophages.

One of the most frequently utilized myeloid reporter mice are Cx3cr1^GFP^, Cx3cr1^Cre^, and Cx3cr1^CreERT2^ knock-in mice ^35,36,38^. Early studies show GFP expression in NK cells and neutrophils from PB in Cx3cr1^GFP^ mice ^36^ but not in Cx3cr1^CreERT2^ and Cx3cr1^Cre^ mice ^38^. However, more recent studies implicate that Cx3cr1^CreERT2^ mice are leaky in brain and can undergo spontaneous recombination even in the absence of TMX ^25,40–43^. The level of spontaneous recombination is markedly higher in Cx3cr1^CreERT2^Ai3^YFP^ mice generated in the Littman laboratory as compared to the ones used in the current study, which were created by the Jung laboratory ^41,43^. Similar reports also show that Cx3cr1^Cre^ mice are non-specific as neurons are positive for the reporter ^42,52^. However, one study identified that the actual reporter mice such as the zsGreen, which has a short distance between the loxP sites may be more susceptible to spontaneous and non-specific recombination as other reporter mice such as the R26-YFP or the R26-CAG-mT/mG, which exhibit markedly less spontaneous recombination ^40^. We show ubiquitous YFP expression in all immune cells from PB and in synovium from Cx3cr1^Cre^Ai3^YFP^ mice. Furthermore, expression is limited to synovial DCs, macrophages and TRMCs in Cx3cr1^GFP^ and Cx3cr1^CreERT2^Ai3^YFP^ mice. Our data are in line with studies by Abram *et al.,* who show reporter expression in neutrophils, basophils, mast cells, NK cells, and lymphocytes using Cx3cr1^Cre^Ai3^YFP^ mice ^44^. In contrast, we did not observe a lack of specificity in PB from Cx3cr1^GFP^ and Cx3cr1^CreERT2^Ai3^YFP^ mice. These data are consistent with studies by Culemann, *et al.,* who utilized Cx3cr1^GFP^ mice to identify the topography of synovial lining macrophages ^5^. Further we show that expression of Cx3cr1 gene and protein are highest in PB CM and NCM as well as CX3CR1^+^ synovial macrophages. Moreover, we previously demonstrated that Cx3cr1^CreERT2^zsGreen mice are not leaky with respect to the synovium^8^. The potential reason for the ubiquitous expression in Cx3cr1^Cre^Ai3^YFP^ mice as opposed to Cx3cr1^GFP^ and Cx3cr1^CreERT2^Ai3^YFP^ mice is that the Cx3cr1 promoter may be activated in early immune progenitors and as a result the YFP becomes expressed ^53^. In contrast, GFP will only be expressed in mice where Cx3cr1 is active even though there might be some GFP that failed to be degraded over time ^36^. Similarly, since we added TMX to Cx3cr1^CreERT2^Ai3^YFP^ mice, we only observe downstream effects of the activated Cx3cr1 promoter. These data demonstrate that Cx3cr1^CreERT2^ mice are more specific for monocyte and macrophage populations except for the brain.

Differential expressions of Cx3cr1 and Ccr2 are commonly analyzed to separate PB CM and NCM^35^. Ccr2^GFP/+^ mice have been used to distinguish tissue-resident macrophages from infiltrating monocytes or monocyte-derived macrophages ^54,55^. However, even though Ccr2 gene expression peaks in CM we observed GFP expression in all PB monocytes. These data are consistent with the concept that NCM are derived from CM ^38^ and thus retain GFP in NCM. The expression of GFP in MHCII^+^ synovial macrophages and a subpopulation of TRMC would indicate that this population may be derived from circulating monocytes as previously reported ^6,8^. These data are also consistent with the reports that loss of CCR2 sequesters monocytes in the bone marrow ^56^. Nr4a1, like CCR2 in CM, controls the differentiation of CM to NCM ^57^. Since we did not observe any RFP expression in synovial macrophages from Nr4a1^RFP^ mice even though Nr4a1 gene is expressed in macrophages these data suggest that the transgene used to generate the Nr4a1^RFP^ mice may be lacking the critical domains i.e. super enhancers necessary for expression in monocytes and macrophages ^58^. Taken together, these data support the notion that a subpopulation of synovial macrophages and TRMCs are derived from CCR2^+^ but not Nr4a1^+^ monocytes.

The lysozyme M cre knock-in (LysM^Cre^) mouse is the 1^st^ Cre recombinase expressing mouse to target myeloid cells including monocytes and granulocytes ^59^ as well as osteoclasts ^60–64^. Since LysM^Cre^ mouse is a knock-in, mice expressing both alleles of Cre are viable, have no abnormalities and respond to innate immune challenge ^65,66^. While LysM^Cre^ mice are not monocyte/macrophage specific as it also effectively deletes floxed DNA sites in neutrophils with the same efficiency as monocytes/macrophages ^59^. Our data also reveal expression of YFP in PB and synovial non-myeloid cells from LysM^Cre/+^Ai3^YFP^ mice consistent with a study that showed a subpopulation of hematopoietic stem cells in LysM^Cre^ YFP mice exhibit Cre activity resulting in reporter expression in lymphocytes ^67^. Additionally, Abram *et al.,* observed YFP expression in splenic lymphocytes, albeit at low levels in LysM^Cre/+^Rosa^EYFP^ mice ^44^. Further studies show LysM promoter activity in peripheral and central neurons ^68^ as well as type II alveolar cells ^65^. We and others utilized the LysM^Cre^ mouse to delete Fas, Bim, Flip, Pd-1, Vamp3, Padi4, Tfr2, Rgs12, Wdfy3, or Sirt1 or express chimeric DR5 genes in myeloid cells ^9–23^. However, in our studies we observed minimal promiscuous deletion in non-myeloid cells. Since we only examine one type of reporter for each marker, other reporter mice have a greater distance in length between the loxP sites, which may yield differences in expression patterns ^9–11,15,69–72^.

The MacGreen transgenic mouse (cfms-EGFP) was the 2^nd^ myeloid reporter mouse generated and expresses EGFP driven by the 7.2 kb promoter of Csf1r/c-fms ^73^. Additional transgenic Csf1r reporter mice were created that utilize the same promoter but different reporter genes such as mApple and yet suffered the same outcome as the MacGreen mouse since ectopic expression was observed in granulocytes, B-lymphocytes and megakaryocytes ^74^. The MacBlue mouse also utilizes the 7.2 kb promoter of Csf1r/c-fms with one exception, it lacks the 180 bp segment containing the trophoblast specific Csf1r promoter, which results in a markedly reduced expression and percent fluorescent positivity in granulocytes as well as tissue macrophages ^75,76^. The Csf1r^Cre^ mouse used in this study was generated using a bacterial artificial chromosome containing all the enhancer and promoter elements of the Csf1r gene ^77^. However, a notable caveat with Csf1r^Cre^ mice is that it also targets neutrophils, lymphocytes, NK cells and monocytes/macrophages in lung and blood ^45^. Our data are consistent with the studies by McCubbey *et al*., as we observed ubiquitous expression of YFP in all immune cells from PB and the synovium ^45^. Grabert *et al* overcame the promiscuous nature of Csf1r expression by generating a Csf1r reporter mice that expresses FusionRed in frame with the protein. Thus, cells that express Csf1r gene but not the protein will be FusionRed negative which includes neutrophils, megakaryocytes and B-cells ^78^. Future studies that can generate a Cre expressing mouse with the same protein stability and expression pattern as CSF1R would greatly move the field of macrophage biology. CD68 is also a known quintessential marker for monocytes and macrophages in humans, while the mouse ortholog of is less specific for myeloid cells ^79^. Thus, the creation of transgenic mice using the 2.9 kb promoter of human CD68 upstream from GFP, rtTA, or CreERT2 (hCD68^GFP^, hCD68^rtTA^ and hCD68^CreERT2^) allows for lineage tracing ^46,80,81^. However, GFP is not only detectable in monocytes and macrophages but also in dendritic cells, neutrophils (Fig 3G, Supplemental Fig 2G) and lymphocytes in PB and lung ^45^, thus limiting hCD68^GFP^ or hCD68^rtTA^ potential for being a macrophage selective reporter mouse ^46^. In contrast, although not examined in our study, hCD68^CreERT2^ mice appear to be more specific for PB NCM and tissue macrophages ^80^.

Over the past years several myeloid reporter mice such as Pf4^Cre^, hMRP8^Cre-ires-GFP^ and Ms4a3^Cre^ were initially thought to be specific for megakaryocytes and/or neutrophils ^82–84^. Notably, Pf4^Cre^ reporter mice were shown to have limited activity in granulocytes from PB but absent in spleen and bone marrow ^44^. However, further studies revealed that Pf4^Cre^ reporter mice have broader expression in a subpopulation of PB monocytes ^44^, interstitial lung macrophages ^85^, splenic macrophages, F4/80^+^ cells in intestine ^49^, hematopoietic stem cells ^86^, lymphoid and myeloid cells responding to inflammatory stimuli ^49,86^, and border associated macrophages, perivascular, pial, dural, IBA1^+^ macrophages but not microglial cells in the brain^25^. Our data show a more specific role for Pf4^iCre^Ai3^YFP^ as we did not observe any YFP in PB cells, while the vast majority of synovial macrophages and half of the TRMC are positive for YFP. These data suggest that Pf4^Cre^ is an ideal reporter for synovial macrophages without affecting synovial lymphocytes or neutrophils. Even though macrophage subsets exhibit variability in Pf4 gene expression, we observed consistently high levels of YFP in Pf4^iCre^Ai3^YFP^ mice. Expression of Pf4 reporter gene in all synovial macrophage subpopulations and TRMC, but not peripheral blood monocytes, indicates Pf4 is related to a maturation process.

Similar to Pf4^iCre^, hMRP8 reporter mice are highly active in granulocytes but marginally present in BAL macrophages and splenic marginal zone macrophages ^44^. Additional studies showed that the same hMRP8 promoter used to generate the hMRP8^Cre/ires-GFP^ mouse is active in monocytes ^84^. In this study, we detected YFP in PB neutrophils and a subset of PB monocytes as well as in NCM from the synovium consistent with previous studies. Nonetheless, the use of hMRP8^Cre-ires-GFP^ mice for arthritis studies is not recommended as the integration of the transgene results in a deletion of 44 kb from chromosome 5 including the loss of Serpine and partial deletion of Ap1s1 ^87,88^. More recently, Ms4a3 gene was identified in granulocyte-monocyte progenitors and subsequently expressed in neutrophils, PB monocytes and tissue macrophages replenished from BM but not in lymphocytes or DCs ^47^. Consistent with Liu *et al.*, we detected YFP primarily in PB and synovial monocytes and granulocytes as well as in tissue resident synovial MHCII^+^ macrophages and TRMC subpopulations derived from PB monocytes in Ms4a3^Cre^Ai3^YFP^ mice even though monocytes do not express Ms4a3 gene. However, we also observed a minor subpopulation of DCs that are YFP in Ms4a3^Cre^Ai3^YFP^ mice. These data suggest that the YFP expression in monocytes as well as DCs are from progenitor cells in Ms4a3^Cre^Ai3^YFP^ mice ^30^.

P2ry12^CreERT2^, Tmem119^GFP^ and Tmem119 ^CreERT2^ mice were generated as microglial specific reporter mice ^24,25^. While P2ry12^CreERT2^Ai3^YFP^ and Tmem119^GFP^ mice do not display any YFP/GFP in circulating cells, a subset of MHCII^-^ synovial macrophages and TRMC are positive in P2ry12^CreERT2^Ai3^YFP^ but not Tmem117^GFP^ mice ^41^. Moreover, expression of P2ry12 was detected in TRMC and in CX3CR1^+^MHCII^-^ synovial macrophages. These data are consistent with studies that show recombination in splenic macrophage subpopulation from P2ry12^CreERT2^ but not Tmem119 ^CreERT2^ reporter mice ^41^.

There are few to no studies that examined the utility of Retnla^Cre^, CD74td^Tomato^, Cd163^iCre^ ^89^ and Tim4d^Cre^ as reporter mice at a global level even though these are known to be macrophage associated genes ^2^. Initially, Lyve1^Cre^ mice were used to target lymphatic endothelial cells ^90^ as well as endothelial cells in high endothelial venules^91^ and then shown to be expressed in hematopoietic cells in fetal liver ^92^. More recently, Lyve-1 reporter mice denote a subset of macrophages, mainly perivascular macrophages ^26–29^. Our comparative analysis between Lyve1^Cre-GFP/+^ and Lyve1^Cre-GFP/+^Ai3^YFP^ mouse models indicate that YFP expression in PB CD11b^-^, granulocytes, DC, CM, and NCM from Lyve1^Cre-GFP/+^Ai3^YFP^ maybe a result from Cre recombinase activity at an early stage in development. We observed expression differences between GFP^+^ and YFP^+^ in Lyve1^Cre-GFP/+^ Lyve1^Cre-GFP/+^Ai3^YFP^ mice, which may be due to the half-life of GFP, a transcriptional efficacy in expression of Cre vs GFP as a result of the IRES (Internal Ribosome Entry Site) ^90^ or that once the stop codon is removed in Ai3^YFP^ mice, there is continuous expression of YFP unlike GFP which is strictly dependent of activation of Lyve-1 promoter. Similar to Lyve-1, CD163 expression is associated with macrophages who monitor the vasculature ^93^. Thus, it is possible that the YFP^+^ populations from Lyve1^Cre-GFP/+^Ai3^YFP^ and Cd163^Cre^Ai3^YFP^ mice are overlapping with each other.

Collectively, this study provides a fate-mapping model atlas for synovial myeloid cells, with a specific focus on macrophages and monocyte-lineage cells. While for some promoters we measured the effectiveness and cell type specificity in the expression level of Cre and CreERT2, spontaneous recombination without TMX, there are limitations as we did not examine the impact of inter loxP distances using different reporter mice, which was examined in microglial cells ^41–43^ and did not determine expression of each promoter in different tissue i.e. skin, spleen, liver, lung, heart, brain. We utilized two CITE-seq datasets to identify genes that are uniquely expressed in each individual macrophage subpopulation and TRMC but not in monocytes. Future studies that generate new transgenic mice for reporter mice should take advantage of published or publicly available scRNA-seq, CITE-seq, or bulk RNA-seq datasets as well as cell atlases. These studies will be crucial for investigations into the tracing of synovial macrophages and monocyte-lineage subpopulations during steady and inflammation.

## Acknowledgments

We would like to thank the Northwestern University Lurie Cancer Center Flow Cytometry Core Facility, supported by NCI Cancer Center Support Grant P30 CA060553. We also thank Dr. Cole Harrington for the gift of Cd74^tdTomato^ mice, Dr. Toby Lawrence for the gift of Cd163^iCre^ mice, Dr. David Voehringer for the gift of Retnla^Cre^ mice, and Dr. John Grainger and Dr. Tovah N. Shaw for the gift of Timd4^Cre^Rosa^YFP^ mice, which were generated by the Genome Editing Unit at The University of Manchester and supported by Senior Fellowship awarded by The Kennedy Trust for Rheumatology Research (John R. Grainger) and BBSRC grant BB/S01103X/2 (Tovah N. Shaw). This research was supported in part through the computational resources and staff contributions provided for the Quest high performance computing facility at Northwestern University and computational resources and staff contributions provided by the Genomics Compute Cluster which are both jointly supported by the Feinberg School of Medicine, the Center for Genetic Medicine, and Feinberg’s Department of Biochemistry and Molecular Genetics, the Office of the Provost, the Office for Research, and Northwestern Information Technology. The Genomics Compute Cluster is part of Quest, Northwestern University’s high performance computing facility, with the purpose to advance research in genomics. The study was supported by R01 AI163742 (Deborah R. Winter), R21 AR080351 (Deborah R. Winter), R01 AR080513 (Harris Perlman & Deborah Winter), and R01 AR075423 (Harris Perlman).

## Materials and Software

### Antibody

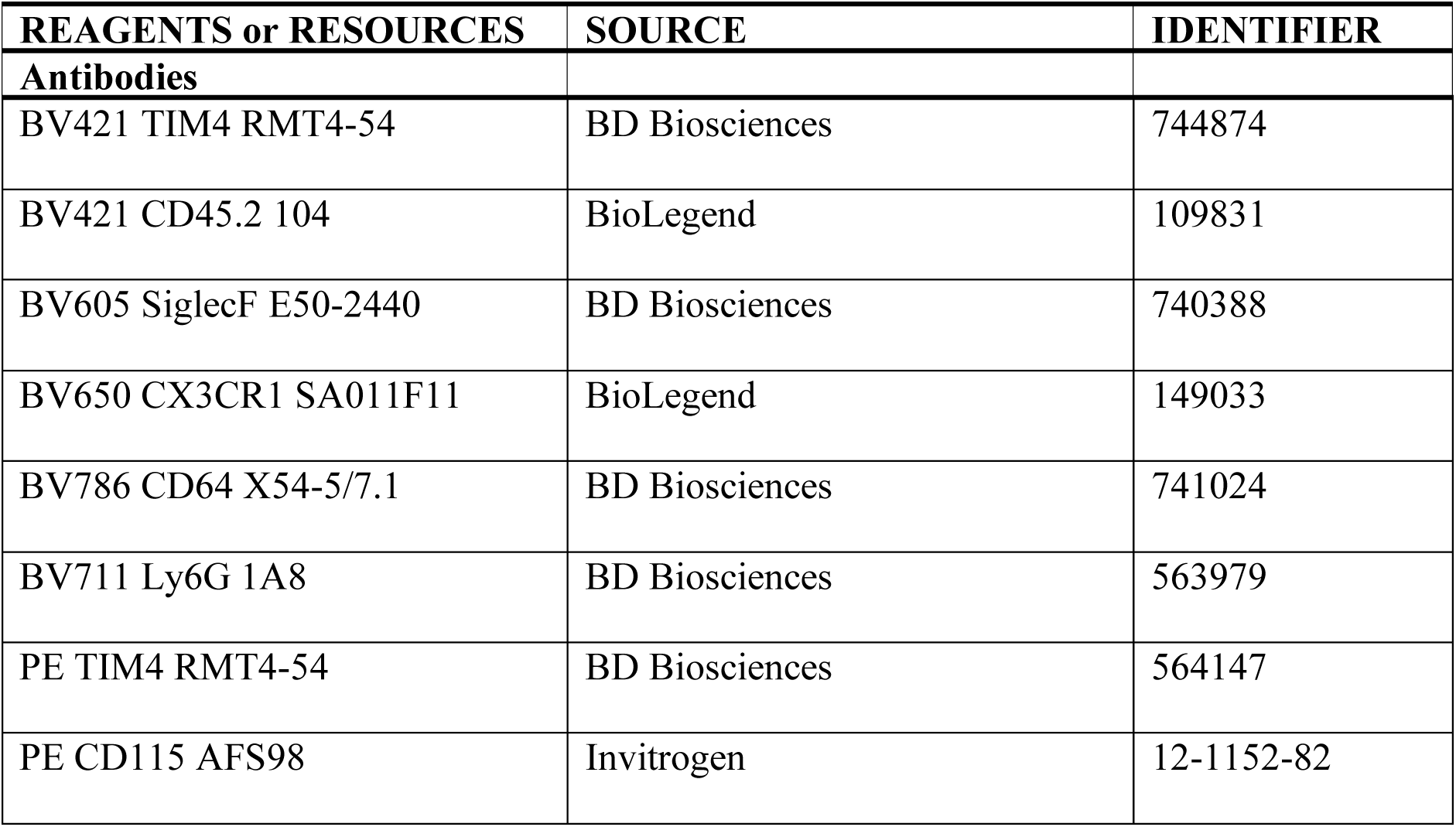
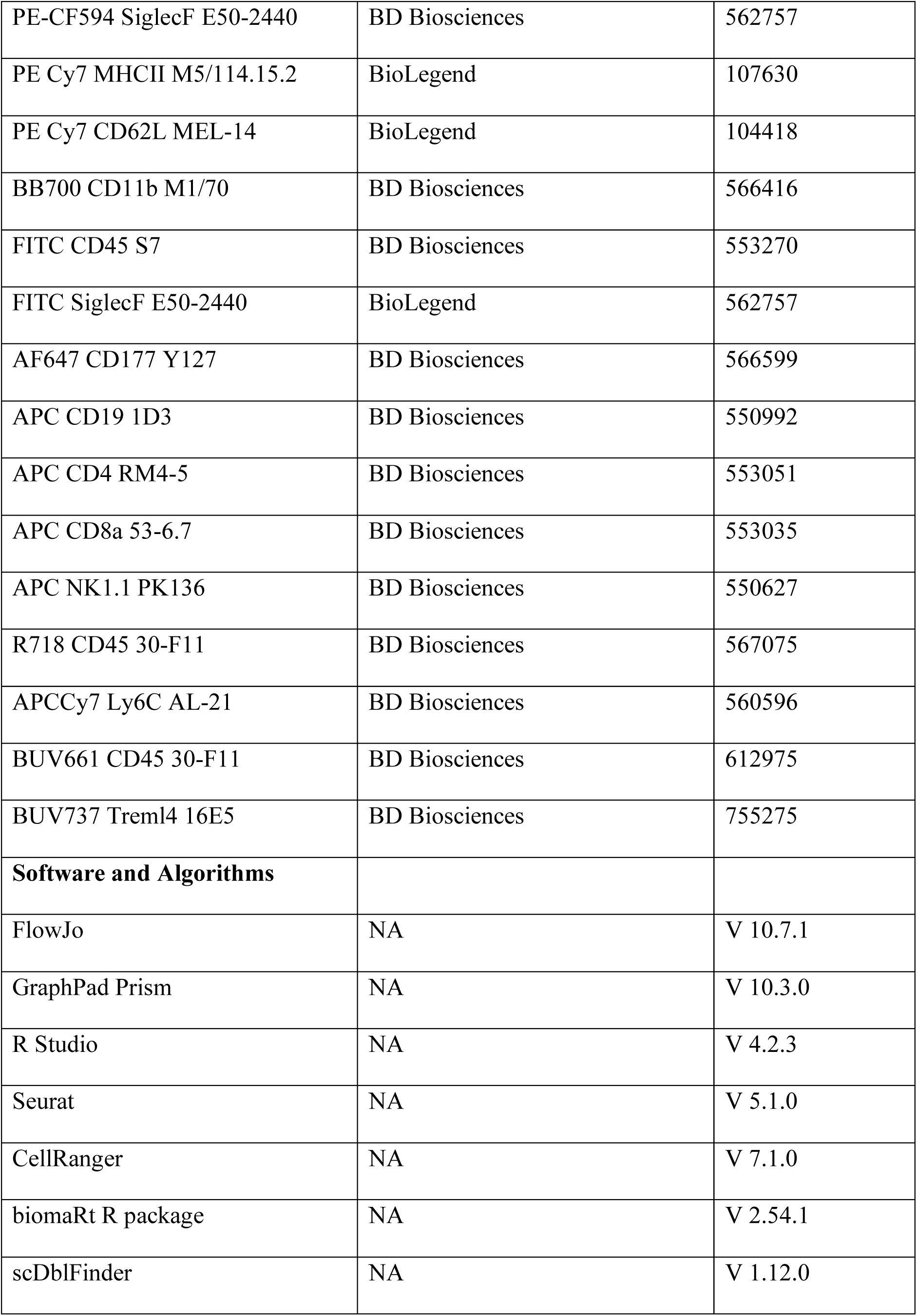
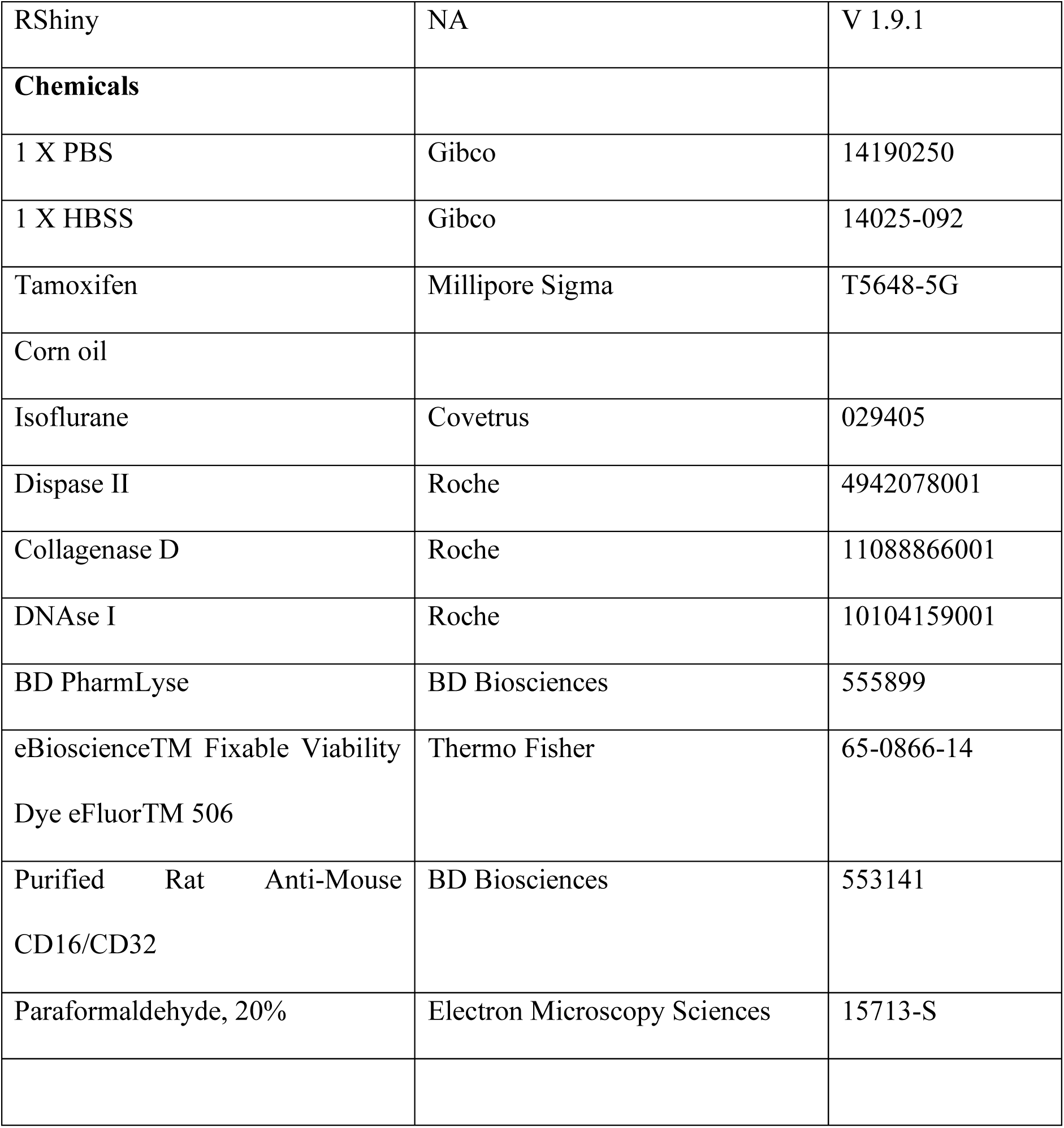

---

**Supplemental Figure 1.**
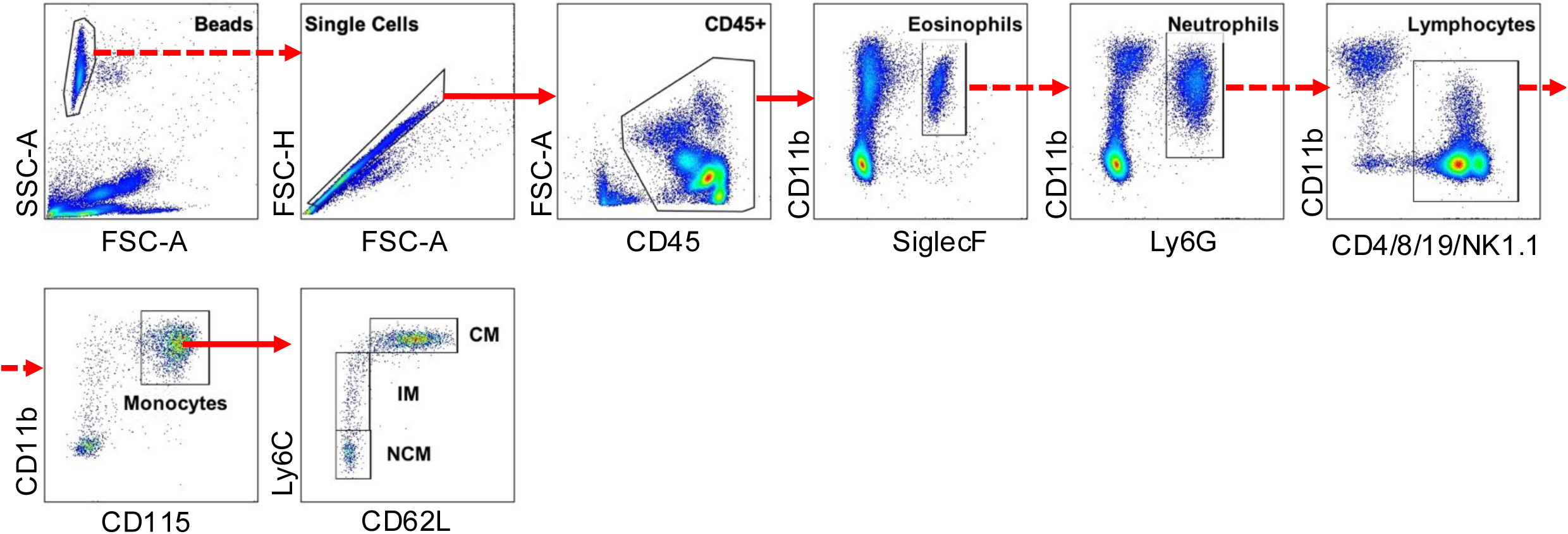
Flow cytometry gating strategy of cells in mouse peripheral blood. Mouse peripheral blood gating strategy for the identification of eosinophils, neutrophils, CM, IM (intermediate monocytes), and NCM. Solid arrows indicate positive gating, while dashed arrows indicate negative gating.

**Supplemental Figure 2.**
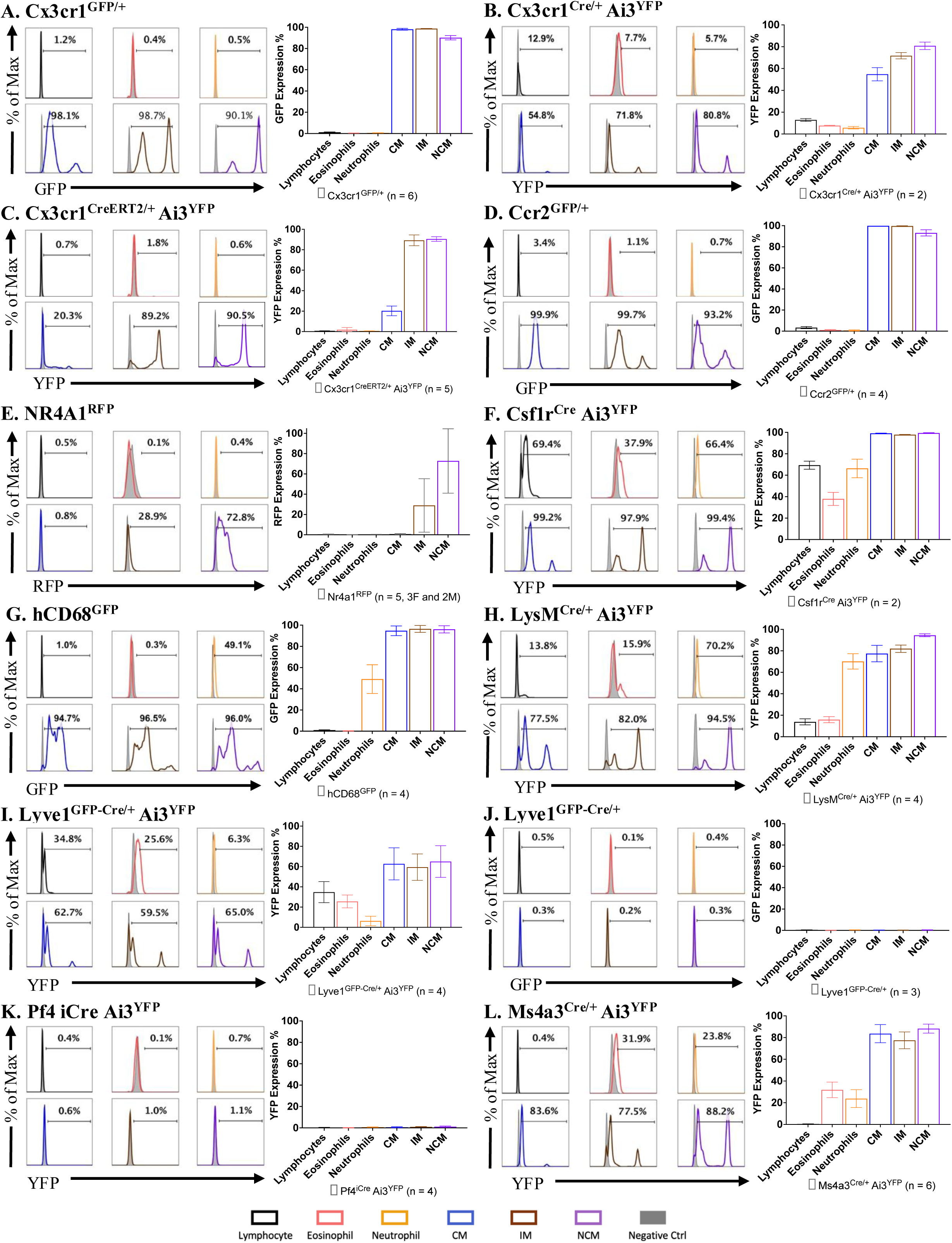
Quantitation of reporter mice expression in PB. Histogram of a representative mouse (percent indicates mean reporter gene-positive across all mice) and bar graph (mean ± SD) summarizing frequencies of reporter gene-positive PB cells in twelve reporter and fate-mapping models. (A) Cx3cr1^GFP/+^ (n = 6; C57BL/6 mice were used as negative controls). (B) Cx3cr1^Cre/+^Ai3^YFP^ (n = 2; Ai3^YFP^ mice were used as negative controls). (C) Cx3cr1^CreERT2/+^Ai3^YFP^ (n = 5; Ai3^YFP^ or Cx3cr1^CreERT2/+^ mice were used as negative controls; 50 mg/kg TAM in corn oil was administered intraperitoneally on day -2 and day -1, and tissues were collected and processed on day 0). (D) Ccr2^GFP/+^ (n = 4; C57BL/6 mice were used as negative controls). (E) NR4A1^RFP^ (n = 5, 3 female and 2 males; C57BL/6 mice were used as negative controls). (F) Csf1r^Cre^Ai3^YFP^ (n = 2; Ai3^YFP^ mice were used as negative controls). (G) hCD68^GFP^ (n = 4; C57BL/6 mice were used as negative controls). (H) LysM^Cre/+^Ai3^YFP^ (n = 4; Ai3^YFP^ mice were used as negative controls). (I) Lyve1^GFP-Cre/+^Ai3^YFP^ (n = 4; Ai3^YFP^ mice were used as negative controls). (J) Lyve1^GFP-Cre/+^ (n = 3; C57BL/6 mice were used as negative controls). (K) Pf4^iCre^Ai3^YFP^ (n = 4; Ai3^YFP^ mice were used as negative controls). (L) Ms4a3^Cre/+^Ai3^YFP^ (n = 6; Ai3^YFP^ mice were used as negative controls).

**Supplemental Figure 3.**
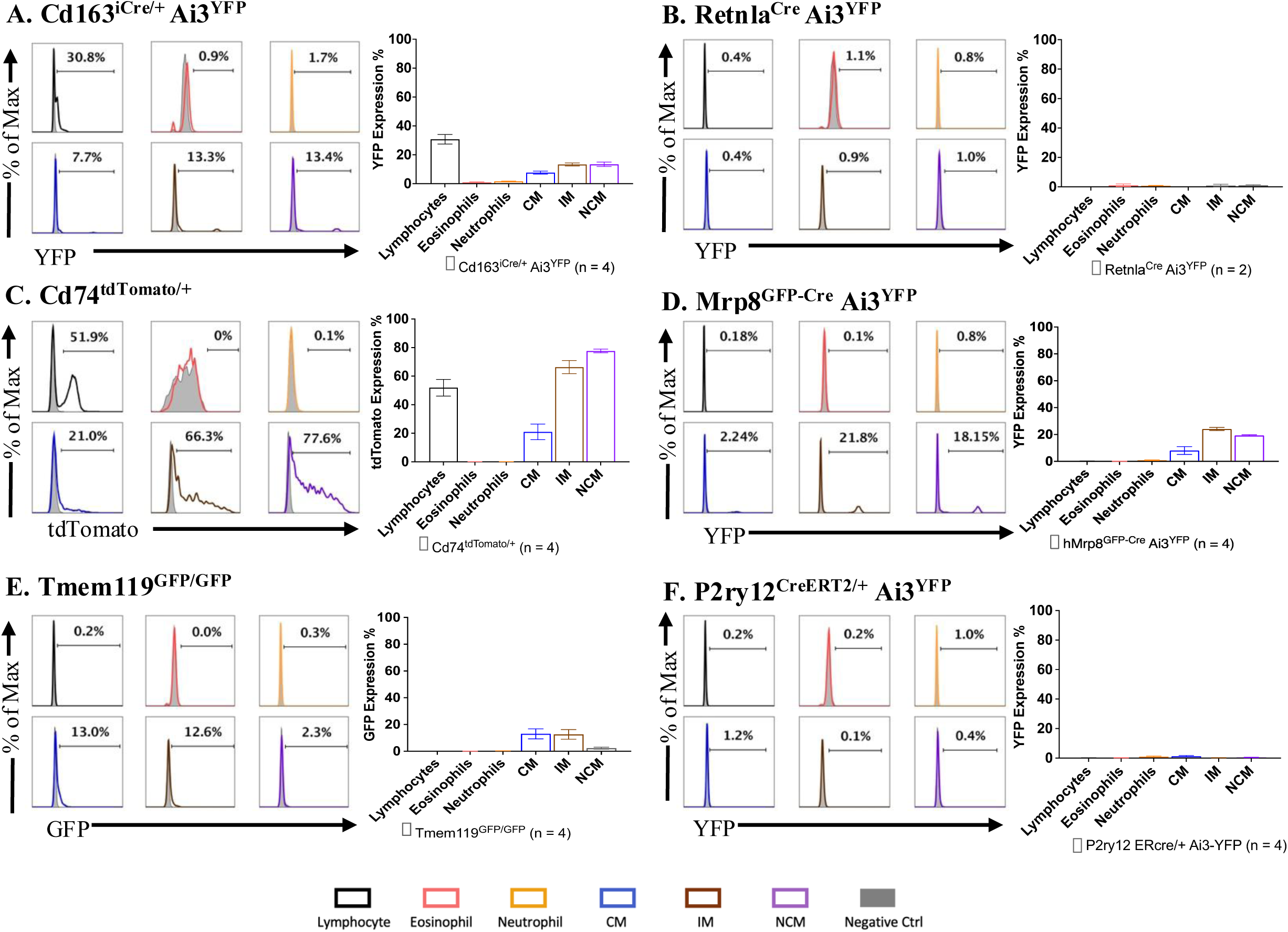
Quantitation of reporter mice expression in PB. Histogram of a representative mouse (percent indicates mean reporter gene-positive across all mice) and bar graph (mean ± SD) summarizing frequencies of reporter gene-positive PB cells in six reporter and fate-mapping models. (A) Cd163^Cre/+^Ai3^YFP^ (n = 4; Ai3^YFP^ mice were used as negative controls). (B) Retnla^iCre^Ai3^YFP^ (n = 2; Ai3^YFP^ mice were used as negative controls). (C) Cd74^tdTomato/+^ (n = 4; C57BL/6 mice were used as negative controls; n = 4). (D) hMrp8^Cre/ires-GFP^Ai3^YFP^ (n = 4; Ai3^YFP^ mice were used as negative controls; blood was withdrawn via heart stick). (E) Tmem119^GFP/GFP^ (n = 4; C57BL/6 mice were used as negative controls). (F) P2ry12^CreERT2/+^Ai3^YFP^ (n = 4; C57BL/6 mice were used as negative controls; 50 mg/kg TAM in corn oil was administered intraperitoneally on day -2 and day -1, and tissues were collected and processed on day 0).

**Supplemental Figure 4.**
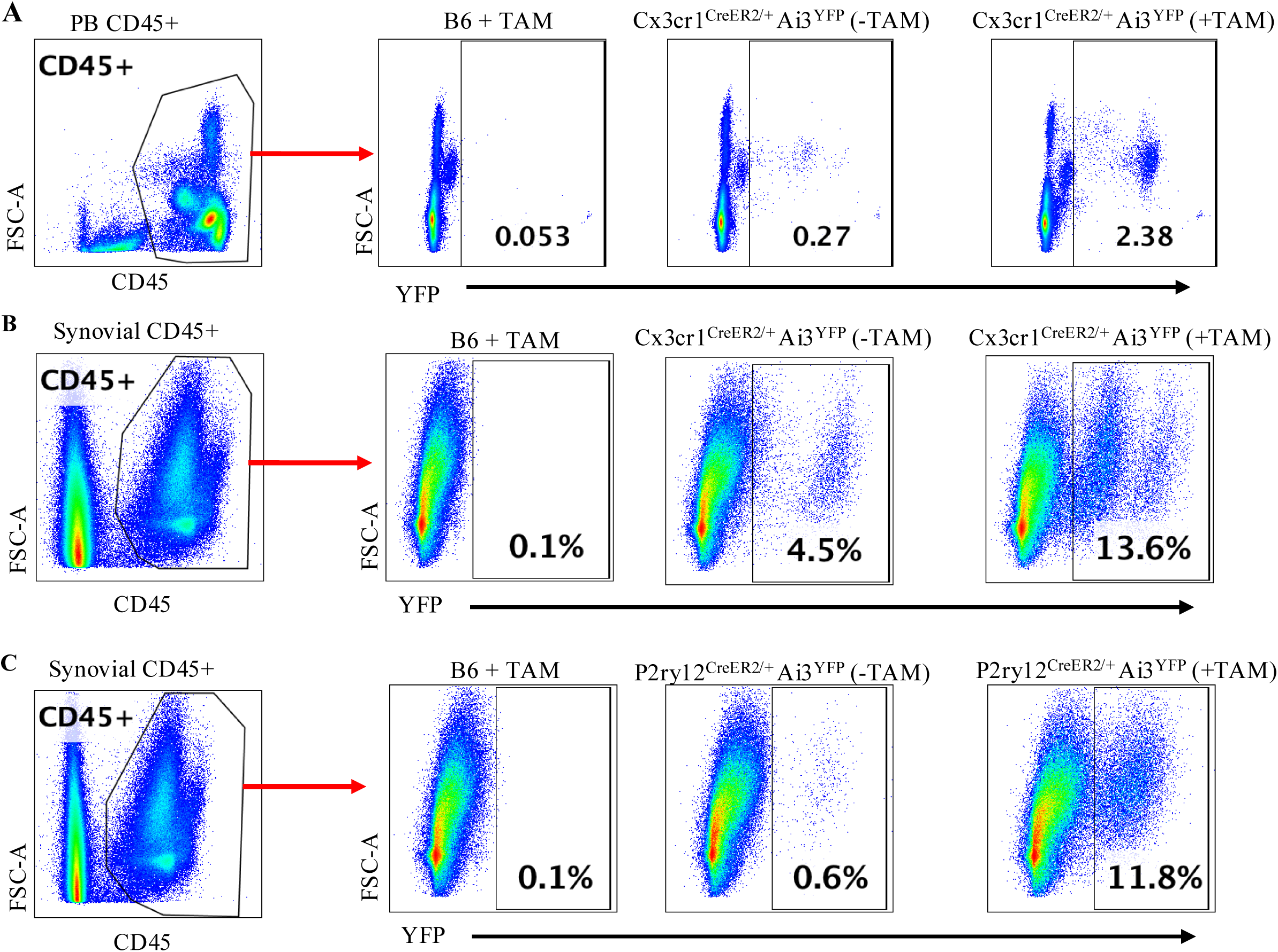
Tamoxifen-independent excision in Cx3cr1^CreER2/+^Ai3^YFP^ and P2ry12^CreER2/+^Ai3^YFP^ mice. (A) YFP expression in CD45^+^ cells in PB from C57BL/6 mouse with TAM, Cx3cr1^CreER2/+^Ai3^YFP^ without TAM, and Cx3cr1^CreER2/+^Ai3^YFP^ with TAM. (B) YFP expression in CD45^+^ cells in synovial tissue from C57BL/6 mouse with TAM, Cx3cr1^CreER2/+^Ai3^YFP^ without TAM, and Cx3cr1^CreER2/+^Ai3^YFP^ with TAM. (C) YFP expression in CD45^+^ cells in synovial tissue from C57BL/6 mouse with TAM, P2ry12^CreER2/+^Ai3^YFP^ without TAM, and P2ry12^CreER2/+^Ai3^YFP^ with TAM. Solid arrows indicate positive gating. TAM at 50 mg/kg in corn oil was administered intraperitoneally on day -2 and day -1, and tissues were collected and processed on day 0.

**Supplemental Figure 5.**
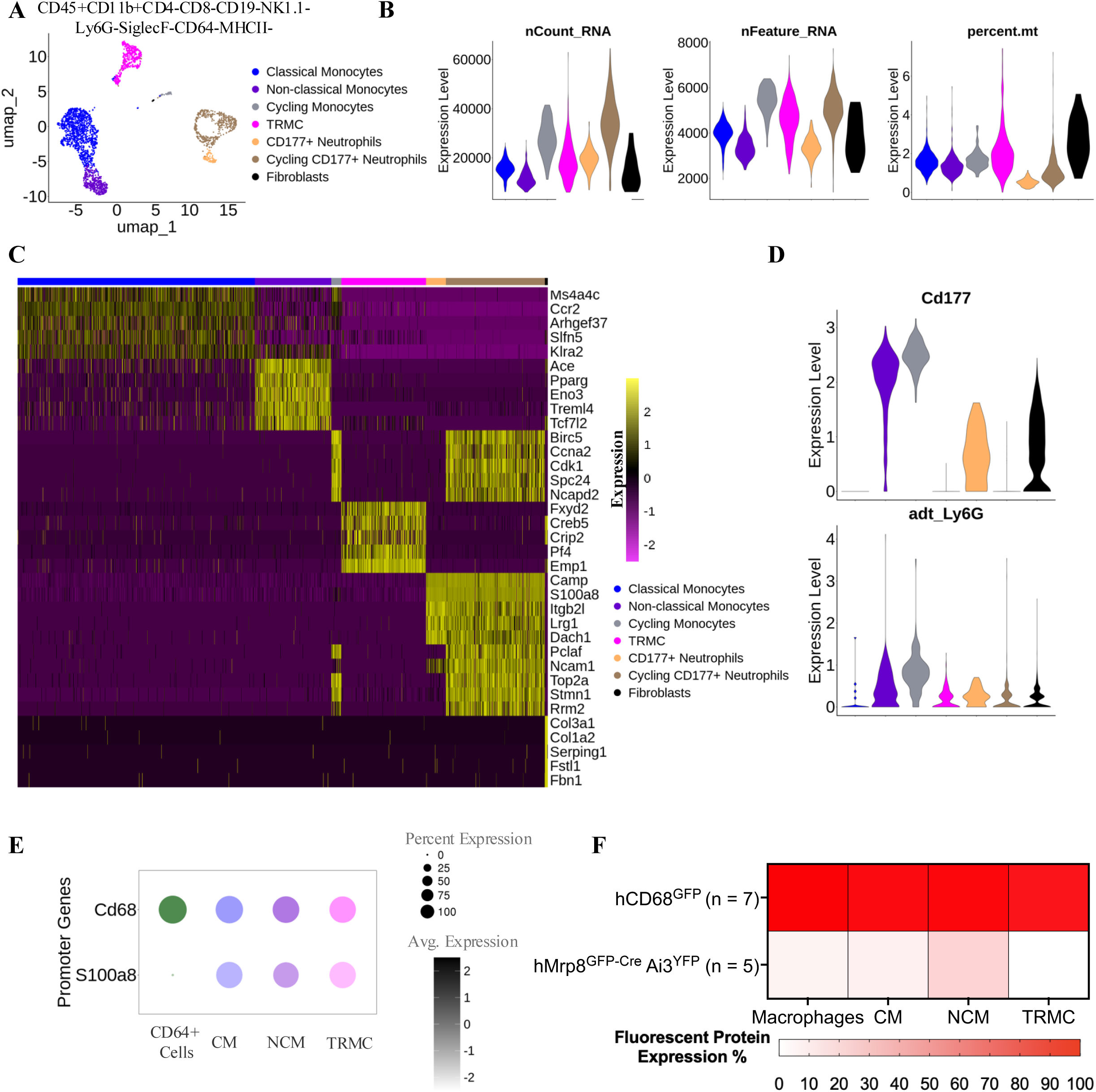
Identification of monocyte-lineage clusters. (A) UMAP depicting seven cell populations of C57BL/6 CD45^+^CD11b^+^CD4^-^CD8^-^CD19^-^NK1.1^-^Ly6G^-^SiglecF^-^CD64^-^ MHCII^-^ cells. (B) Quality controls of the seven cell populations showing the numbers of nCount_RNA, nFeature_RNA, and percent of mitochondrial genes (percent.mt) of C57BL/6 CD45^+^CD11b^+^CD4^-^CD8^-^CD19^-^NK1.1^-^Ly6G^-^SiglecF^-^CD64^-^MHCII^-^ cells. (C) Heatmap of selected five differentially expressed genes (DEG) of the seven cell populations identified in A. (D) Feature plots of the gene expression of Cd177 and ADT expression of Ly6G in the cell types identified in A. (E) Bubble plots showing the average expression of the Cd68 and S100a8 of CD64^+^ cells (macrophages), classical monocytes (CM), non-classical monocytes (NCM), and TRMC. (F) Heatmap of the percent of cells with detectable fluorescent protein in hCD68^GFP^ and hMrp8^GFP-Cre^Ai3^YFP^ across macrophages, CM, NCM, and TRMC defined by flow cytometry.

**Supplemental Figure 6.**
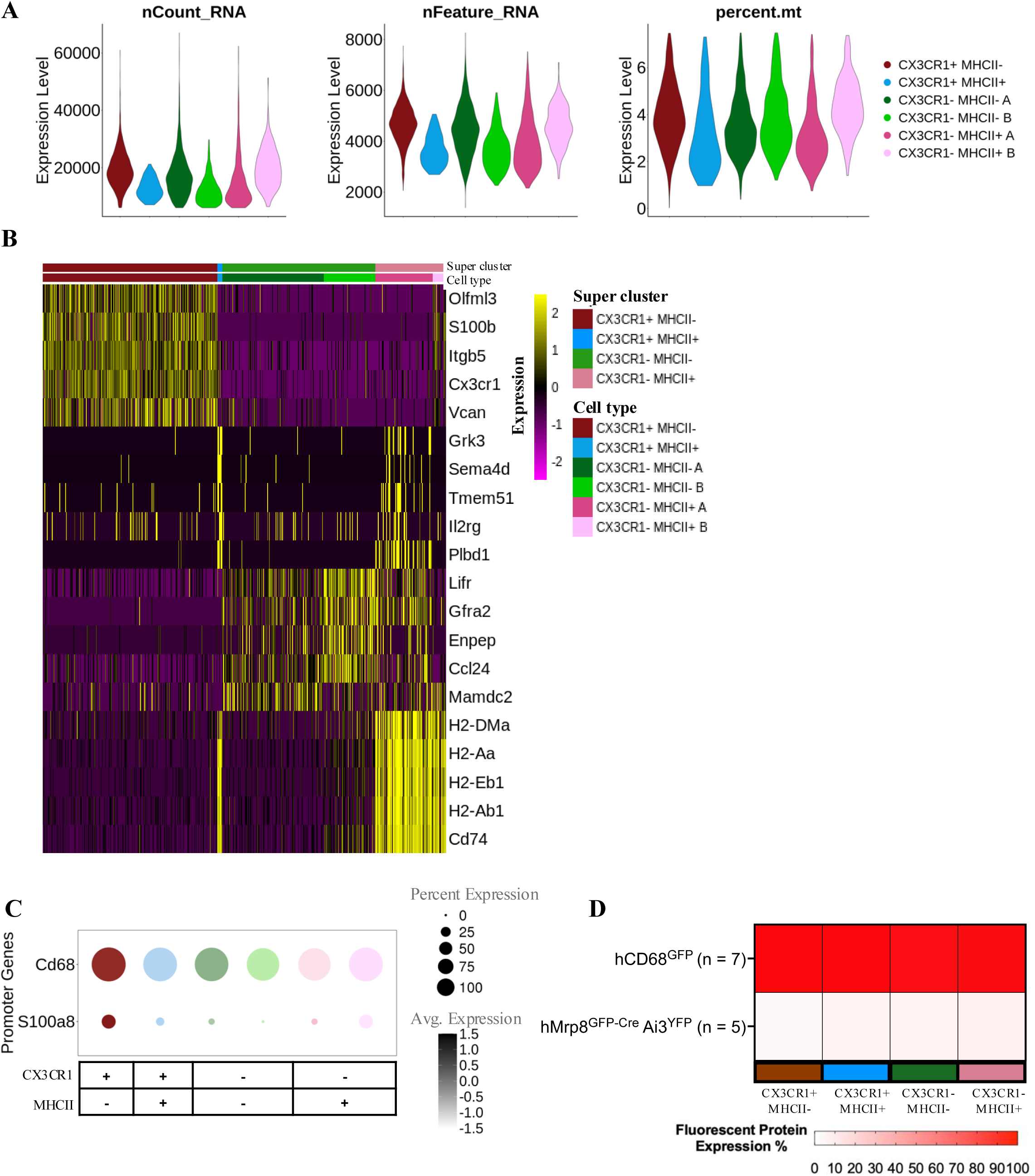
Identification of synovial macrophage clusters. (A) Quality controls of the six macrophage subsets from CD64^+^ dataset showing the numbers of nCount_RNA, nFeature_RNA, and percent of percent.mt of C57BL/6 CD45^+^CD11b^+^CD4^-^CD8^-^CD19^-^NK1.1^-^ Ly6G^-^SiglecF^-^CD64^+^ cells. (B) Heatmap of five DEG for the four super-clustered synovial macrophage subsets. (C) Bubble plots showing the average expression of the Cd68 and S100a8 of the macrophage subsets identified in Figure 5A. (D) Heatmap of the percent of cells with detectable fluorescent protein in hCD68^GFP^ and hMrp8^GFP-Cre^Ai3^YFP^ across the four macrophage subsets defined by flow cytometry.

